# Meta-analysis of the autism gut microbiome identifies factors influencing study discrepancies and machine learning classification

**DOI:** 10.1101/2022.03.18.484910

**Authors:** Aries Chavira, Eric Hou-Jen Wang, Robert H. Mills

**Author notes:** **Corresponding Author:** Robert H. Mills, Precidiag Inc, 313 Pleasant Street, Watertown, MA 02472, USA.

## Abstract

Autism Spectrum Disorder (ASD) is a severe neurodevelopmental disorder and accumulating evidence has suggested that dysbiosis of the gut microbiome plays an essential role. However, a body of research has investigated the ASD gut microbiome without consensus as to whether or how the ASD microbiome differs from neurotypical children. Here, we evaluate the underlying factors leading to study discrepancies by performing a meta-analysis on reprocessed 16S ribosomal RNA gene amplicon (16S) sequencing data. We compile a total of 1,740 samples across 13 carefully selected published studies together with samples from the American Gut Project, and analyze the data in aggregate and from a per-study perspective. We observed increased *Bifidobacterium*, Actinobacteria, and *Prevotella* among ASD individuals across cohorts. We further identified associations to Desulfovibrionales, Deltaproteobacteria and *Prevotella* that were dependent upon which 16S variable regions were sequenced. Utilizing machine learning (ML), we obtained increased accuracy in ASD classification using data collected from certain territories, on younger subjects, on unrelated age-matched rather than related controls, on samples with increased sequencing depth and when accounting for sex differences. Our work provides compelling evidence that the gut microbiome is altered in ASD patients, and highlights novel factors that are important considerations for future studies.

## Introduction

Autism Spectrum Disorder (ASD) represents a group of neurodevelopmental disorders characterized by a spectrum of behavioral, speech, and social impairments. The CDC recently reported that diagnoses have increased to 1 in 44 children (2.3%) in the United States^1^. Autism is also included in the Global Burden of Disease (GBD), highlighting ASD as a disease of increased global importance^2^. The precise etiology of autism is unknown but is thought to be multifactorial. Research into a genetic basis for autism has identified combinations of genes that are linked to ASD, however genetics are estimated to only contribute to 10-30% of ASD cases^3; 4^. Given that gastrointestinal issues and immune system dysregulation are prevalent comorbidities of individuals with autism, accumulating studies have investigated whether the complex consortium of microbes in the gut is involved in the etiology and/or pathogenesis of ASD^5-8^.

The human gut microbiota has emerged as an important factor in a large range of diseases including emerging links to neurological conditions including ASD. Studies have reported ASD-like behaviors in mice transplanted with the feces from ASD children, which suggests that the microbiome is causally involved in the development of ASD^9; 10^. Importantly, small trials of fecal microbiota transplants in humans have reported improved behavioral outcomes^11-13^. Mechanistically, there are multiple hypotheses for how gut microbes may be involved in ASD including the maternal immune activation model^14^, the production of neurotransmitters such as serotonin^15^ or GABA^16^, modulating vagus nerve signaling^17^, and production of molecules such as 4-ethylphenylsulfate^8^ or short-chain fatty acids^18^.

There are now over 250 scientific articles or reviews discussing the microbiome and ASD. The results of individual studies assessing whether there are alterations in the gut microbiome of human populations with ASD have drawn different conclusions as to the strength of an association between ASD and controls^19-22^. However, systematic reviews and meta-analyses conclude that individuals with ASD indeed have a significantly altered microbiome^23; 24^. In some accounts, large shifts in the ecology of the microbiome is observed, including significant separation in beta-diversity between ASD and controls as well as differences in broad taxonomic categories such as the ratios of the Firmicutes to Bacteroidetes phyla^20^. In contrast, others report very little changes in these metrics, no change^22^, or a significant shift in the opposite direction^25^. Further, a recent report found very few ASD-associated microbes, and showed that a less diverse diet correlated with ASD status which the authors suggest may be responsible for differences in the ASD microbiome^19^.

Here we sought to identify the underlying factors influencing the conflicting results within studies investigating the ASD microbiome. We hypothesized that some between-study variability could be explained by study-specific demographics of the population being studied. For example, aspects such as the geographic location and age of participants could influence the results of a study given that these factors are known to influence the microbiome^26; 27^. Additionally, research has shown how differences in 16S sequencing methods can impact the microbial profiling of individuals with ASD^28^. Accordingly, we systematically reprocessed and reanalyzed existing ASD microbiome data with a focus on how study design variables impacted our results. We evaluated the effect that 16S rRNA sequencing methods, sex, geographic location, control type and age had on detecting differences in the ASD microbiome as well as how these factors may influence the accuracy of predicting ASD status via machine learning models. Our data is derived from thirteen carefully selected published case-control cohorts, and one curated cohort from the American Gut Project (AGP)^29^. We analyzed the datasets both in conglomerate and individually to gain more insight on how the ASD gut microbiome differs from developmentally neurotypical individuals.

## Results

### Included Studies and Sample Demographics

Following the Preferred Reporting Items for Systematic reviews and Meta-Analyses (PRISMA)^30^, we curated a dataset consisting of fourteen different case control cohorts^20-22; 29; 31-40^ (Fig. 1a). The resulting dataset comprised a total of 1,740 samples, of which 888 were diagnosed with ASD and 852 were typically developing controls. Our largest cohort was manually curated from participants of the AGP (n=524), with other large cohorts from Dan et al. (n=286)^20^ and Chen et al. (n=246, Fig. 1b)^32^. Control subjects were selected from the AGP in a manner strictly controlling for technical variability while finding best-matches based on a variety of selected variables. This approach resulted in a control cohort that largely reflected the demographics of the AGP’s ASD subjects (Extended Data Fig. 1). In total, a majority of samples were from males, which follows the general trend of increased incidence of ASD amongst boys (Fig. 1c). The studies typically included participants between the ages of 2-6 years old, with notable exceptions being Kong et al. with an age range of 6-62 and Kang et al. of 7-16 (Fig. 1d)^31; 33^. Thirteen of our cohorts represent samples from the United States (n = 1079) and China (n = 866), and one study, Zurita et al., 2019 was conducted in Ecuador (n = 50)^36^ (Fig. 1e). In addition, a large portion of samples (n = 1280) were processed targeting the 16S rRNA V4 hypervariable region, while 357 samples were sequenced from the 16S V3-V4 region and only 103 were processed from the V1-V2 region. For controls, ten studies utilized an age and sex matched sample from unrelated controls, two studies utilized siblings, and two others included the mothers of ASD children (Fig. 1f). More details on the demographics of subjects from each study can be found in Extended Data Table 1.

**Figure 1.**
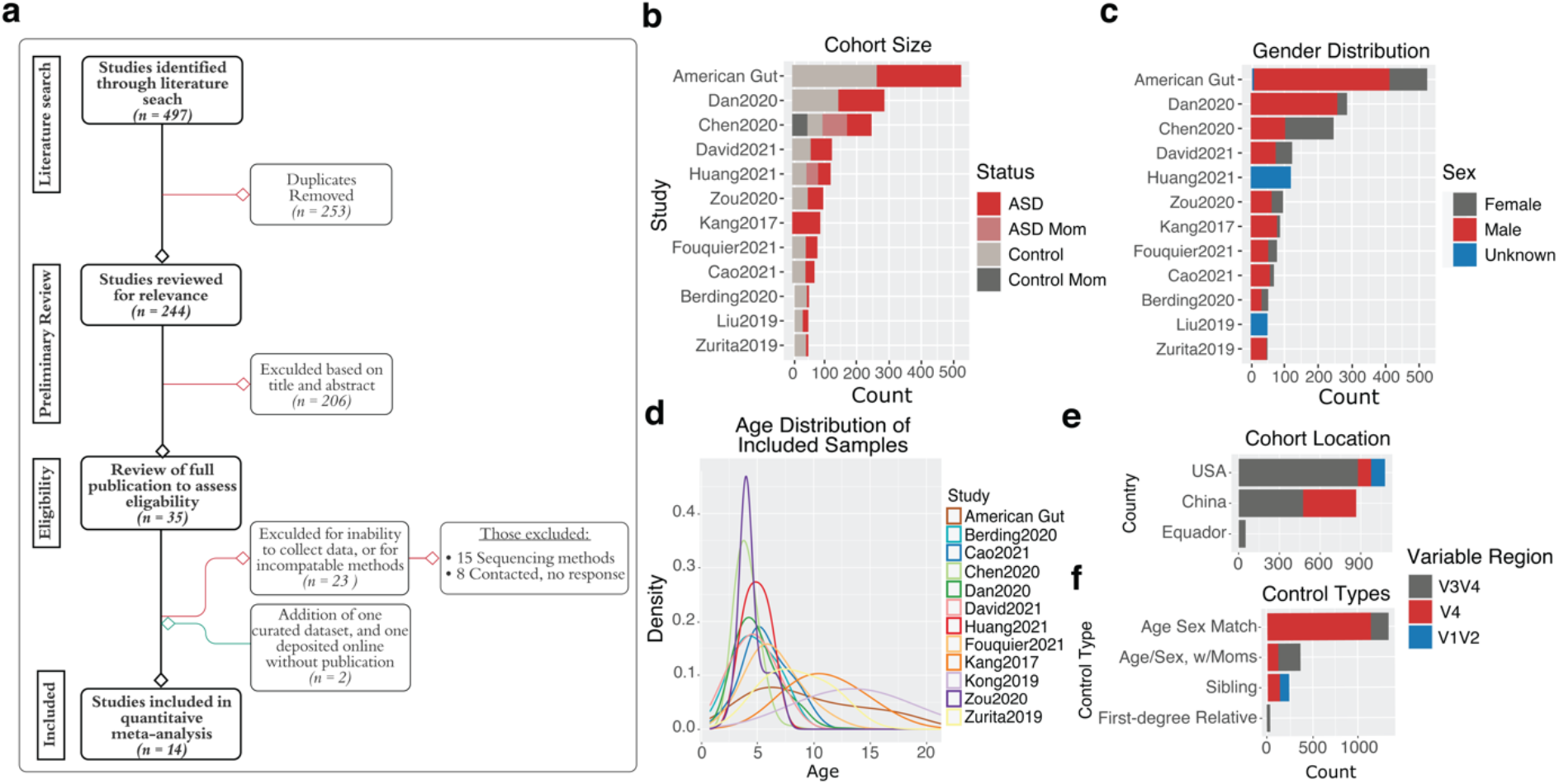
PRISMA flow chart and cohort demographics. **a**, Flow chart of studies assessed for eligibility to be included in the final analysis. Thirty-five studies were reviewed in full to assess inclusion eligibility. A total of 15 were included in the final analysis with a total sample size of n = 1740 (ASD = 888, controls = 852). **b**, Total sample sizes for each cohort plotted in decreasing order and colored by ASD or control status. **c**, Total sample sizes of each cohort plotted in decreasing order colored by sex. **d**, Density plot of age for each cohort. **e**, Samples broken down by country in decreasing order, and colored by 16S hypervariable region. **f**, Samples stratified by control type in decreasing order and colored by 16S hypervariable region.

### Evaluating differences in ASD microbiome diversity

To address the question of whether microbiome diversity is altered in ASD, we assessed multiple alpha and beta-diversity metrics from both a per-study and an aggregated dataset approach. For beta-diversity, we tested two commonly utilized distance metrics - unweighted and weighted UniFrac - which are phylogenetically-informed metrics^41^. In the aggregate, we observe a significant clustering by study for both the unweighted and weighted UniFrac distance metrics, though the effect was notably stronger for the unweighted UniFrac distance (unweighted UniFrac PERMANOVA *P* < 0.001, pseudo-F = 42.15; weighted UniFrac PERMANOVA *P* < 0.001, pseudo-F = 18.54) (Fig. 2a-b). This observation suggests that study-unique microbes of low abundance increase study separation, and drive study discrepancies. While less striking than the study effect, we observed significant clustering by ASD status in unweighted UniFrac distance (PERMANOVA *P* < 0.001, pseudo-F = 3.99), but not in weighted UniFrac distance (PERMANOVA *P* = 0.162, pseudo-F = 1.48) (Extended Data Fig. 2a-b). Strong separation was also observed based on which 16S hypervariable region was used during sequencing (Extended Data Fig. 3a), highlighting the importance of accounting for which variable region was used. We also observed differences related to sequencing depth and geographic location in unweighted UniFrac distances (Extended Data Fig. 3b-c), though these results are confounded by the effect of variable region and study.

**Figure 2.**
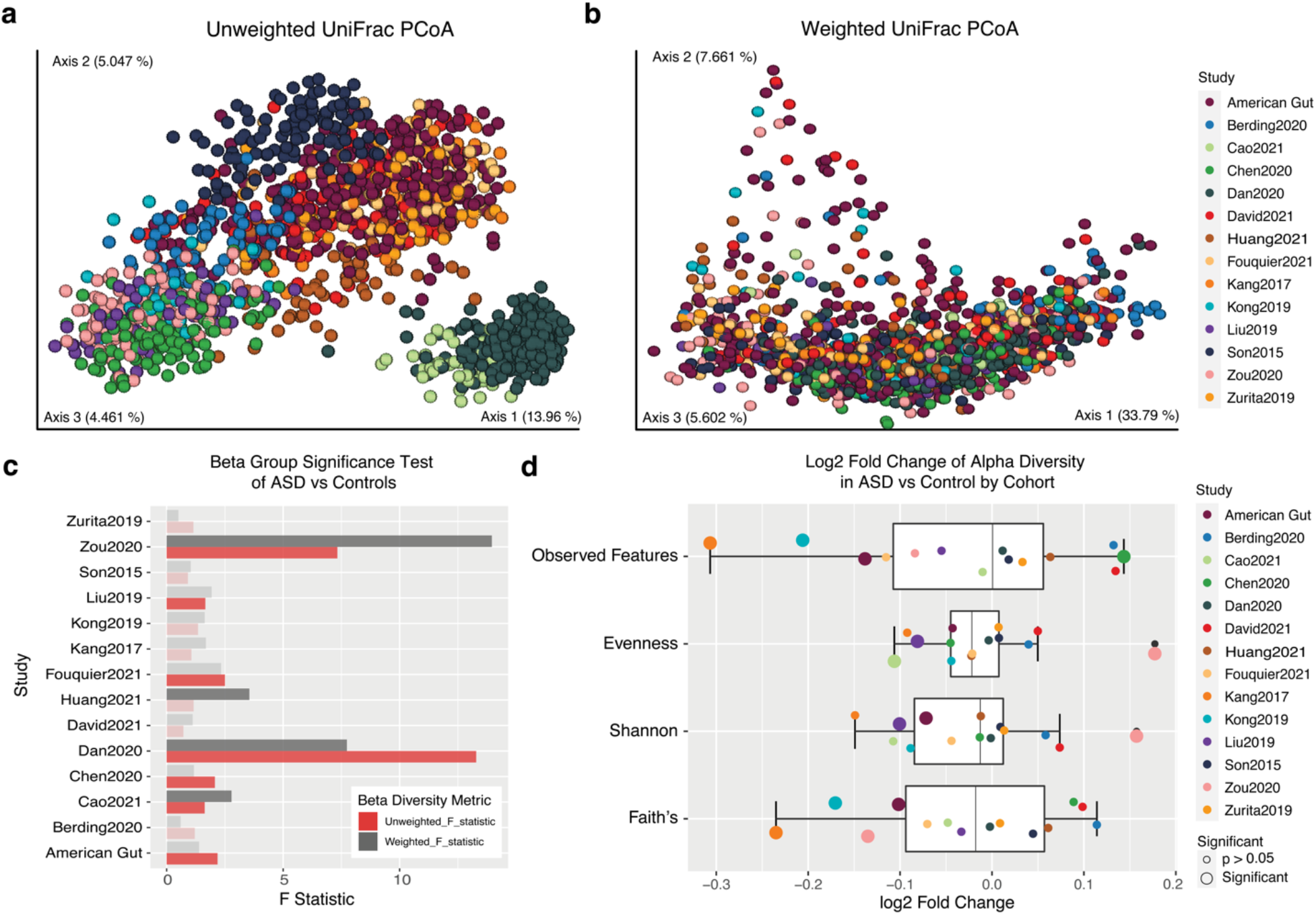
Association of ASD status to microbiome alpha and beta-diversity. **(a-b)** PCoA from the unweighted (a) and weighted (b) UniFrac distance matrix respectively, of samples colored by cohort (n = 1740). **c**, Pseudo-F statistic from a PERMANOVA beta-group significance test of ASD vs control from unweighted and weighted UniFrac distance metrics. Bars with full color opacity refer to PERMANOVA tests with a *P* < 0.05. **d**, Log2 fold change in alpha-diversity values of ASD to controls for each cohort and sized by significance. Boxplots show the median, quartiles and 1.5x inter-quartile range of the data distribution. P-values were generated from unadjusted two-tailed t-tests.

Given that a strong confounding effect on beta-diversity results were driven by the source of study, we further analyzed each cohort individually. Eight of fourteen cohorts showed statistically significant differences (PERMANOVA *P* < 0.05) in the ASD gut microbiome compared to neurotypical controls from either the weighted or unweighted UniFrac distance metrics (Fig. 2c). These results indicate that there is variability in detecting beta-diversity differences between ASD and controls, however, differences can be detected in 57% of studies, as well as when analyzing datasets in aggregate.

We also analyzed each study for the effect of ASD status on within-sample microbial diversity. We utilized four alpha-diversity metrics: Faith’s phylogenetic diversity, Shannon diversity, evenness, and observed features. For all metrics, the study effect was large and required us to analyze each of the fourteen cohorts individually. We show that for each of the four alpha-diversity metrics utilized, there were at least three studies that showed statistically significant differences in the alpha diversity values between ASD and neurotypical controls (Students’ t-test, *P <* 0.05) (Fig. 2d). Of the 14 cohorts, 50% showed statistically significant differences in alpha diversity between ASD and neurotypical controls in at least one of the four metrics (Fig. 2d). In general, the average alpha diversity values for ASD subjects were anywhere from 0.09 to 2.5% lower than controls across datasets, except for observed features where the average values for ASD subjects were 0.04% higher than controls (observed features = +0.04%, evenness = -2.5%, Shannon = -0.09%, and Faith’s = - 1.2%). These results suggest an unclear association between ASD and alpha-diversity.

### Differences in microbial relative abundance between ASD and controls

We next assessed differences in microbial relative abundances between ASD and control subjects at a per-study level, aggregating the data at different phylogenetic levels. We observed that few phyla were consistently significantly different, with the most commonly significant phylum being Bacteroidetes where four studies reached statistical significance (Fig. 3a). Despite not reaching statistical significance, a majority of cohorts showed increased Actinobacteria, and Proteobacteria (Fig. 3a). We also investigated the ratios of specific phyla as there are many reports of altered ratios of Firmicutes/Bacteroidetes among ASD subjects^21; 42; 43^. We found that the ratio of Firmicutes/Bacteroidetes was elevated in ASD individuals in nine cohorts (Fig. 3a). In contrast, the ratio of Firmicutes/Bacteroidetes was decreased in Zou et al., Chen et al., and Kang et al., (Fig. 3a). Following up on reports of significant differences in the Firmicutes/Bacteroidetes ratio between Eastern and Western societies^44; 45^, we tested for differences in this ratio of samples in the USA and China and did not find a significant difference (unadjusted *P* = 0.29) (Extended Data Fig. 4a).

**Figure 3.**
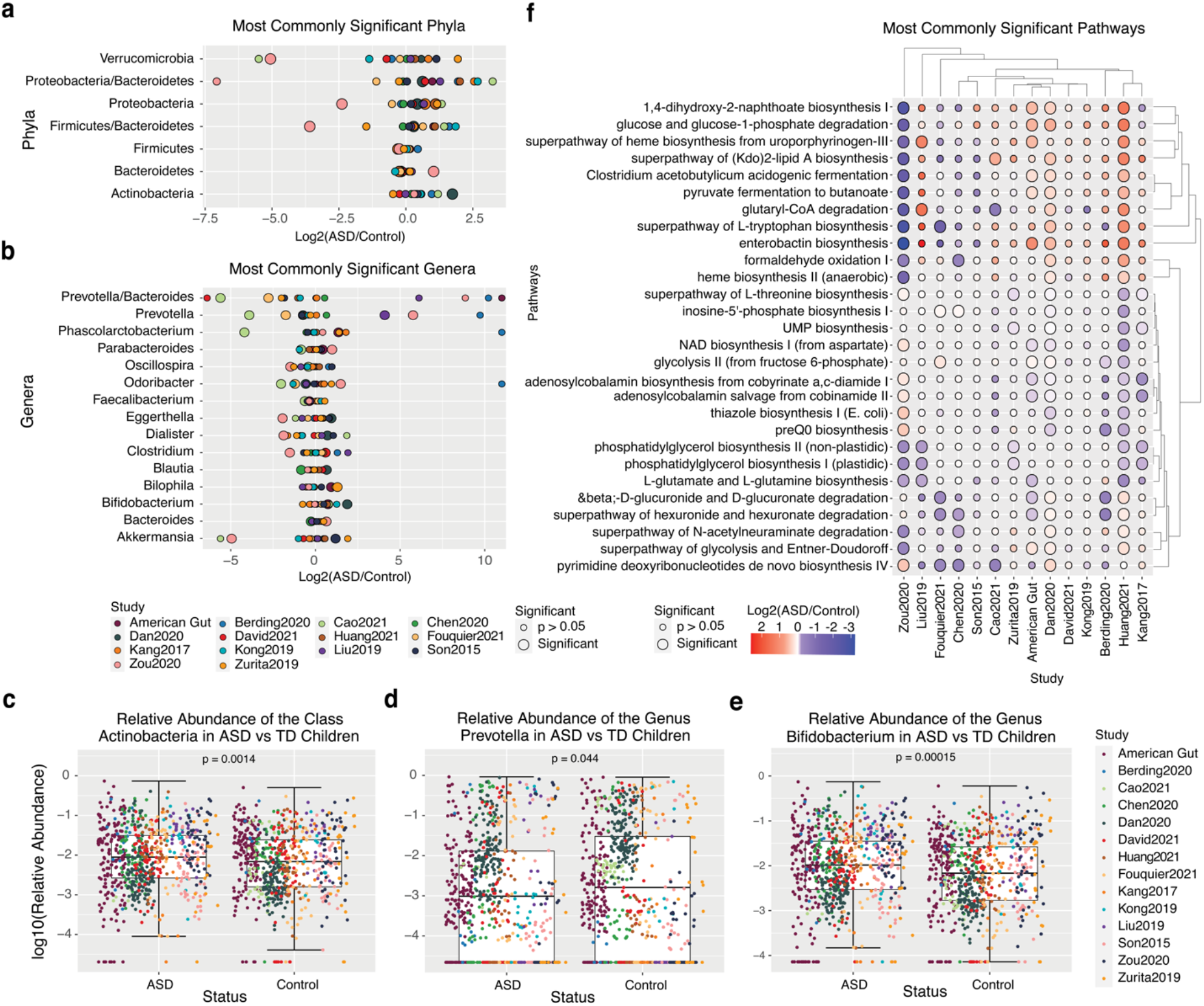
Taxonomic and functional differences between ASD and controls. **(a-b)** Log2 fold change in the relative abundance of ASD/controls of the most commonly significant phylum (a) and genus (b). Individual points are colored by cohort with the size indicating a significant difference between ASD and controls (unadjusted two-tailed t-tests p-value < 0.05). **(c-e)** Per-sample plots of the log10 transformed relative abundance of each sample for the class Actinobacteria (c), the genus *Prevotella* (d), and the genus *Bifidobacterium* (e), stratified by ASD status. P-values from wilcoxon two-tailed t-tests are shown comparing ASD to Control values (Actinobacteria n = 1478, P = 0.0014; *Prevotella* n = 1478, *P* = 0.044; *Bifidobacterium* n = 1478, P = 0.00015). Boxplots show the median, quartiles and 1.5x inter-quartile range of the data distribution. **f**, Commonly, significant microbial pathways when comparing ASD and controls. Plotted are the log2 fold changes of the normalized pathway abundance values in ASD/control for the most commonly significant bacterial metabolic pathways for each cohort processed individually. Hierarchical clustering of studies and pathways was performed using Euclidean distance. Points are colored by log2 fold change and sized by significance (unadjusted two-tailed t-tests *P* < 0.05).

The most common genus to have a significant difference in its relative abundance was *Prevotella*, for which five cohorts reached statistical significance (unadjusted *P* < 0.05) (Fig. 3b). However, these studies were split as to whether *Prevotella* was increased or decreased in ASD subjects (Fig. 3b). Further, features at the class and order level were also inconsistently increased or decreased in ASD, though twelve of our fourteen cohorts showed an increased abundance of the class *Enterobacteriales* among ASD subjects (Extended Data Fig. 4b-c).

When analyzing our data in aggregate, there were also few taxa that generalized to our whole dataset. A few notable exceptions include the class Actinobacteria, which was significantly higher in ASD children compared to controls (Wilcoxon t-test *P* = 0.0014, Fig. 3c). Despite the inconsistent per-study findings for *Prevotella*, we found the general trend indicating a decrease in ASD samples compared to controls (Wilcoxon t-test, *P* = 0.044, Fig. 3d). One of our strongest findings among the aggregate data confirms past associations with increased *Bifidobacterium* among ASD subjects (Wilcoxon t-test, *P* = 0.00015, Fig. 3e). In general, significant taxonomic associations in the ASD microbiome were not generalizable across studies, with the few notable exceptions listed above.

### Chorismate links several microbial pathways commonly altered in ASD

Despite the taxonomic inconsistency across studies, it is possible that the functional pathways encoded by the microbiome of ASD subjects are more generalizable. To test this possibility, we predicted the metabolic pathway abundances of each sample and compared ASD to control subjects. After assessing the most commonly significant pathways, we observed two major clustering patterns of pathways that were generally increased vs generally decreased in ASD subjects (Fig. 3f). Of particular interest, we noted that several of the commonly identified pathways shared a common molecular precursor, chorismate.

Chorismate is a product of the shikimate pathway, which is absent in mammals. Chorismate can be transformed into several important molecules including menaquinols and ubiquinols. We identified 18 pathways involving the biosynthesis of various menaquinols, and ubiquinols that were significant in three or more studies (Extended Data Fig. 3d). These pathways included the biosynthesis of menaquinols 6-13 which were largely increased in ASD apart from Zou et al., which showed a significant decrease among ASD subjects (Extended Data Fig. 3d). Menaquinones and ubiquinones are vital molecules involved in bacterial anaerobic and aerobic respiration respectively and can be synthesized from chorismate. This could indicate that microbial species with differing reliance on anaerobic and aerobic respiration are commonly found in ASD.

Aromatic amino acids (AAA’s) are also products originating from chorismate. From our analysis, L-tryptophan biosynthesis was significantly altered (*P* < 0.05) in four studies and increased among ASD subjects in nine of the fourteen studies (Fig. 3f). However, pathways related to the other aromatic amino acids, L-tyrosine and phenylalanine were not as commonly significant, though L-tyrosine degradation was increased in ASD within eight of our fourteen cohorts (Extended Data Fig. 4e). We also noticed that other amino-acid related pathways, such as L-threonine biosynthesis and L-glutamate and L-glutamine biosynthesis, were significantly decreased in five cohorts each (Fig. 3f). These results support previously noted differences in amino acid metabolism in ASD patients^46; 47^ and expand the analysis to new cohorts.

Other molecules derived from chorismate include siderophores and folate which we also found altered among several ASD studies. Enterobactin is a metal chelating siderophore, which we identified to be significantly altered in Dan et al., Huang et al., and Zou et al., (Extended Data Fig. 4e). We further identified tetrahydrofolate biosynthesis to be significantly decreased in 4 studies (Extended Data Fig. 4e). Further, adenosylcobalamin biosynthesis from cobyrinate (also known as Vitamin B12), was decreased in ten of our fourteen cohorts with four cohorts reaching concordant significance (Fig. 3f). Vitamin B12 catalyzes the conversion of homocysteine to methionine and tetrahydrofolate, playing a rate-limiting step on folate levels^48^.

### Evaluating machine learning performance on ASD status prediction

The relationship between the microbiome and ASD may be non-linear and therefore not easily identified through traditional statistical approaches. Therefore, we next utilized various ML methods to assess the extent to which the ASD gut contains a signature distinguishable from that of neurotypical individuals. We evaluated the performance and accuracy of predicting ASD status using eleven different classification algorithms across the aggregate of our fourteen datasets (Fig. 4a). All eleven models performed at least marginally better than chance as determined by having an area under the receiver operating characteristic curve greater than 0.5 (Fig. 4a). Our top performing models, gradient boosting classifier (GBC) and adaboost classifier (ABC), were able to distinguish ASD individuals from controls with an area under the curve (AUC) greater than 0.6 (GBC; AUC = 0.62 ± 0.03, ABC; AUC = 0.61 ± 0.02) (Fig. 4a). Using the F1 score of each model to assess both precision and recall, we found that GBC and ABC models were performing at 0.62 ± 0.03, and 0.62 ± 0.02 respectively (Extended Data Fig. 5a). Third, we assessed the performance of each model by evaluating each model’s classification accuracy. Both the GBC and ABC models reported the largest percent accuracy of 62% ± 0.03, and 61% ± 0.02 respectively (Extended Data Fig. 5b). Across all the metrics, the GBC performed the best at classifying ASD status and was therefore selected for downstream analyses.

**Figure 4.**
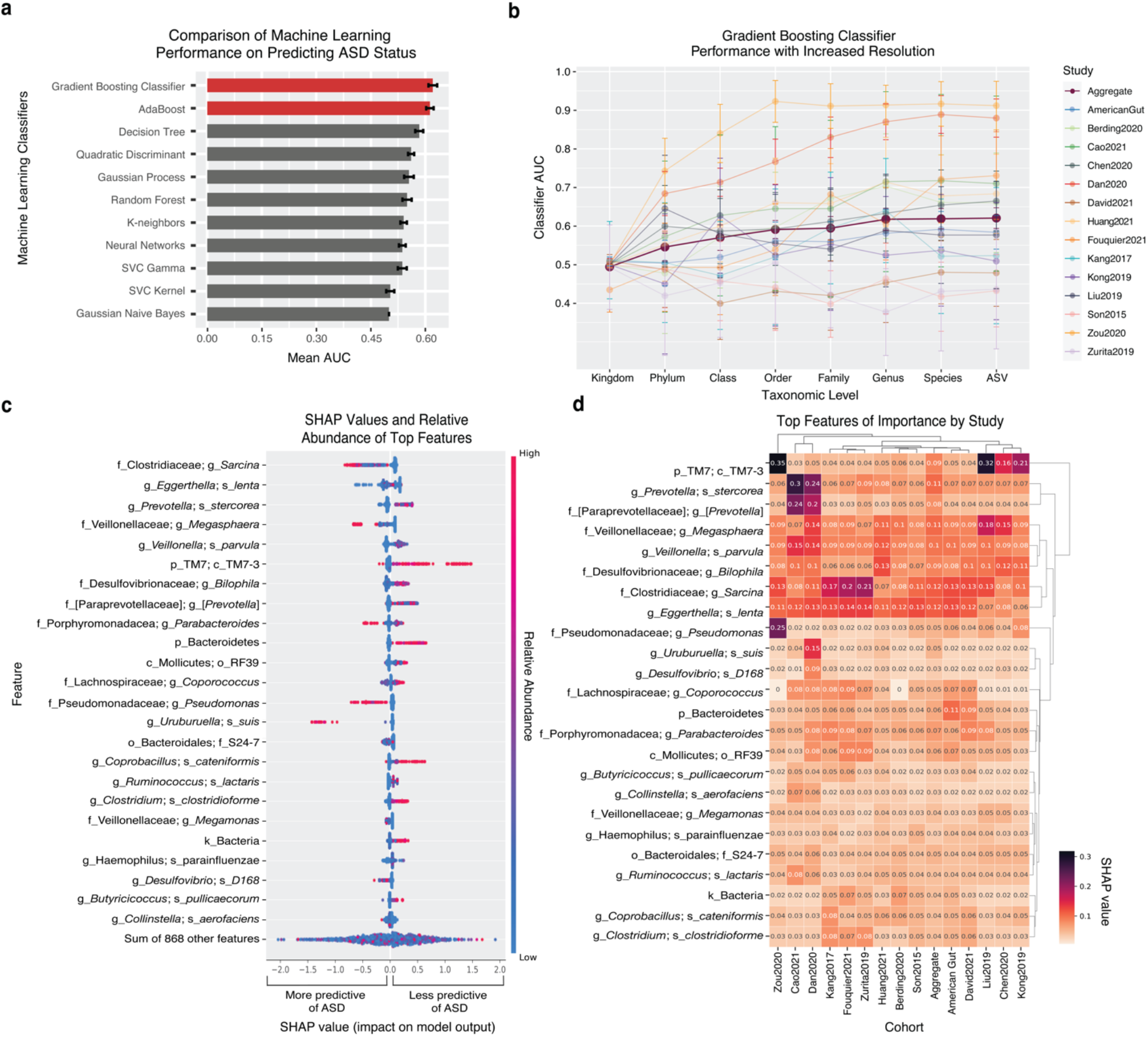
Comparing machine learning performance on ASD status prediction. **a**, Bar plots of the mean AUC +/-SEM from 50 iterations. Eleven classification algorithms were tested using the aggregated dataset of ASV’s given the most resolved taxonomic assignment. Models highlighted in red performed best and were further tested. **b**, The mean AUC of the gradient boosting classifier models trained and tested on both the aggregated data, and each study individually on reads counts collapsed at different taxonomic levels. Error bars show the standard error from 5-fold cross validation repeated 10 times. **c**, The relative abundance and SHAP value for each sample of the top 25 features are plotted in decreasing feature importance from the gradient boosting classifier model trained on the aggregate dataset of ASV’s given the most resolved taxonomic assignment. **d**, Cluster heatmap generated and colored by the SHAP values of the top 25 features from the gradient boosting classifier model trained and tested on the aggregate data and each cohort independently. Clustering of both cohorts and features was performed using hierarchical clustering of Euclidean distances.

As our initial ML model comparison was performed on data aggregated by taxonomic annotation, we were interested in seeing how taxonomic resolution impacted the model’s performance and accuracy. We performed fifty iterations of randomized train-test splits utilizing the GBC model on read count data either un-collapsed or aggregated at seven different taxonomic levels (Kingdom, Phylum, Class, Order, Family, Genus, Species), as well as the highest resolution of the assigned taxonomy. Increasing taxonomic resolution improved the model’s accuracy, with only marginal gains past genus level classification (Fig. 4b). We also observed similar trends when performing the same analysis with both the adaboost and decision tree classifiers (Extended Data Fig. 5c-d). Training and testing each dataset individually, GBC models performed better than chance (AUC > 0.5) in all but three data sets (78%), and five had AUC’s above 0.70 (Fig. 4b). Further, the GBC model of each cohort was correlated with both the unweighted and weighted UniFrac Pseudo-F statistic from the beta groups significance PERMANOVA test for ASD status (Pearson correlation coefficient; weighted UniFrac = 0.79, unweighted UniFrac = 0.75) (Extended Data Fig. 6). These results indicate that the ASD gut microbiome likely contains information that would be beneficial to assessing ASD status.

We next investigated the important features in distinguishing individuals with ASD from controls. Our top three features from the GBC model trained and tested on the aggregated, highest-resolution dataset were amplicon sequencing variants (ASVs) from the genus *Sarcina* (family Clostridiaceae), *Eggerthella lenta*, and *Prevotella stercorea* (Fig. 4c). Among the top 25 features, increased relative abundances of ASVs from *Sarcina*, the genus *Pseudomonas*, and *Uruburuella suis* were important for ASD classification, while increased abundances of ASVs from class TM7-3, the phylum *Bacteroidetes, Coporobacillus cateniformis*, and *Clostridium clostridioforme* were important for control classification (Fig. 4c). To provide more context to our findings, we further assessed the top 25 features from the aggregate dataset for their importance to predicting ASD status within GBC models generated on each individual dataset. Hierarchical clustering of the importance scores of these features to each data set revealed two main groupings (Fig. 4d). Of particular interest was a group of features that contained similarly high importance scores across all datasets. This group included ASVs from *Eggerthella lenta*, the genus *Sarcina*, the genus *Bilophila, Veilonella parvula*, and the genus *Megasphaera*. These top features highlight that despite the heterogeneity of results among studies, there are several microbial features that can consistently be utilized for ASD classification.

### Evaluating the influence of study design on study outcomes

To address the variability in ASD microbiome results reported in the literature along with those highlighted in our work, we sought to evaluate how factors of study design can impact the findings of an ASD microbiome study. Some of these factors include, but are not limited to, 16S sequencing methods, geographic location, sex, age, and the relationship of the control subject to the ASD subjects. Regarding various technical choices of 16S sequencing methods, we evaluated the effect of sequencing depth and choice of 16S hypervariable regions.

Among the 16S hypervariable regions included in our data, one study targeted the V1-V2 region, 5 studies utilized the V3-V4, and 8 studies used the V4 region (Extended Data Table 1). V1-V2 samples were excluded from this analysis given the limited number of studies conducted utilizing that variable region. We found that studies utilizing the V3-V4 variable region had a significantly higher relative abundance of the genus *Prevotella* in ASD subjects (*P* = 4.61E-3) while studies targeting the V4 region had a significantly lower relative abundance *Prevotella* in ASD subjects (*P* = 5.6E-4) (Fig. 5a). This was also consistent when comparing the ratio of ASD/control relative abundances for *Prevotella* from a per-study perspective (*P* = 0.011) (Fig. 5a). We also found the ratio of the genera *Prevotella* to *Bacteroides*, the order *Desulfovibrionales*, and the class *Deltaproteobacteria* were different between the V3-V4 and V4 regions (per-study Wilcoxon t-test *P =* 0.017, 0.011, and 0.011 respectively) (Fig. 5b-d). When analyzing our data in aggregate as opposed to on a per-study basis, we noted that significant differences could be found for *Desulfovibrionales* and *Deltaproteobacteria* among V3-V4 studies, while no difference was found among V4 samples (both decreased in V3-V4 ASD with a *P* = 0.015).

**Figure 5.**
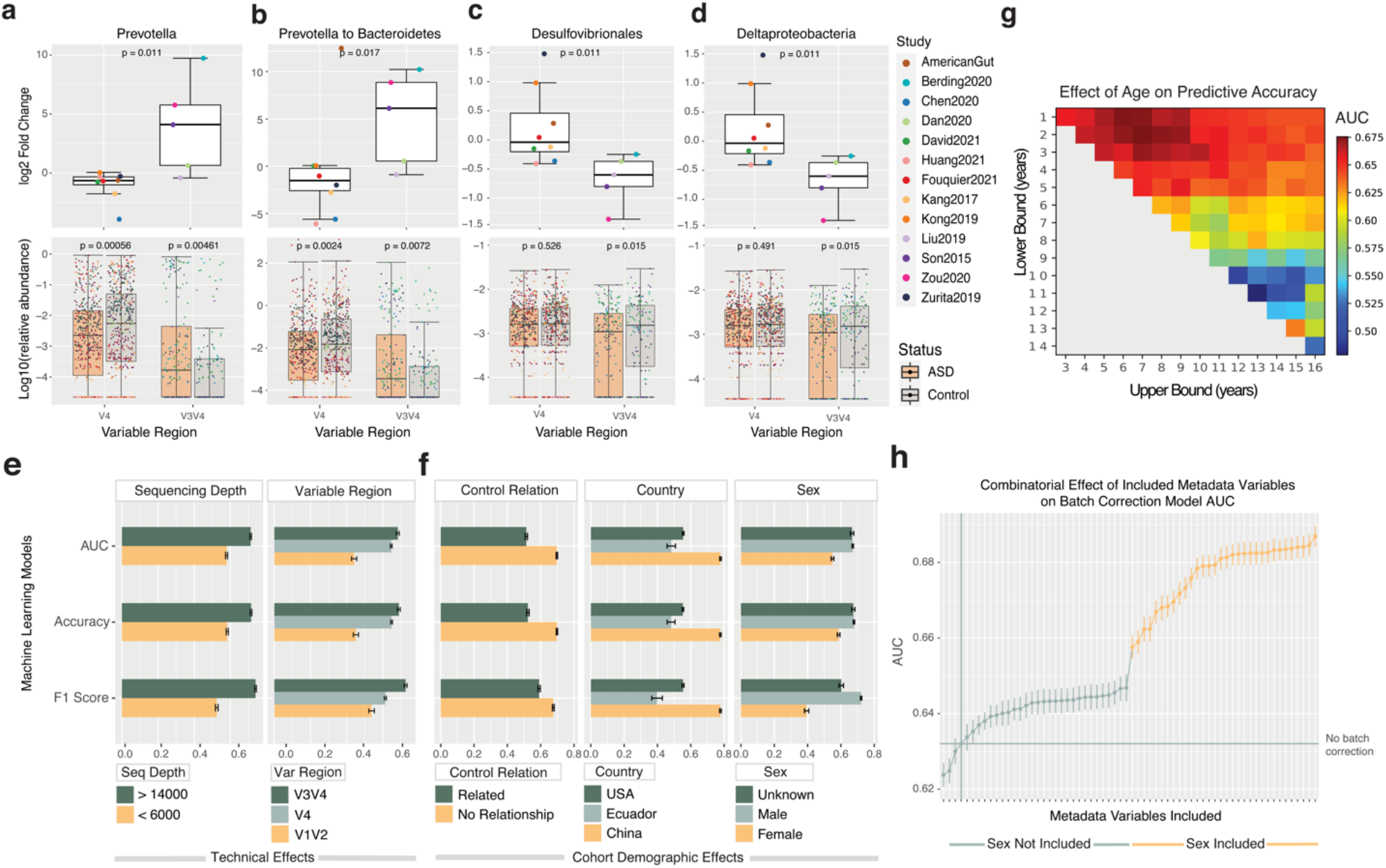
Evaluating study design factors that influence the performance of machine learning models and taxonomic abundance. **(a-d)** Top; Per-study log2 fold change of ASD to controls of the genus *Prevotella*, ratio of *Prevotella*/Bacteroidetes, order Desulfovibrionales, and class Deltaproteobacteria of data sequenced from the V4 and V3-V4 hypervariable regions respectively. Bottom; Log10 transformed relative abundance of the genus *Prevotella*, ratio of Prevotella/Bacteroidetes, order Desulfovibrionales, and class Deltaproteobacteria in each sample from the V4 and V3-V4 regions respectively, split by ASD status (V4 n = 1024, V3-V4 n = 352). P-values calculated by unadjusted Wilcoxon two-tailed t-tests. Boxplots show the median, quartiles and 1.5x inter-quartile range of the data distribution. **e**, Machine learning AUC, accuracy, and F1 statistic of the gradient boosting classifier’s trained to predict ASD status from samples split by sequencing depth, and variable region (sequencing depth; > 14000 n = 851, > 6000 n = 640, variable region; V3-V4 n = 356, V4 n = 1032, V1-V2 n = 103). **f**, AUC, accuracy, and F1 statistic of the gradient boosting classifier at predicting ASD status from samples split by control relationship, country, and sex (control relationship; related controls n = 462, unrelated controls n = 1029, country; USA n = 761, Ecuador n = 46, China n = 684, sex; unknown n = 168, male n = 1021, female n = 302). **g**, Heatmap of the GBC model’s average AUC trained and tested on samples binned at each potential age range from 1-16. (e-f) Barplots show mean +/-SEM. **h**, AUC, and standard error of the mean (SEM) bars from 50 iterations of GBC models trained on the aggregated dataset subjected to batch correction. A reference line is shown representing the mean AUC of baseline model. Models trained on data where sex was, and was not, included as a variable in the batch correction are indicated by color.

Despite identifying discrepancies between taxonomic associations to ASD by sequenced variable region, ML models trained on both data sets performed well above chance while V3-V4 data outperformed V4 data (average AUC of 0.65 and 0.70 for V4 and V3-V4 models respectively) (Fig. 5e). These results suggest that there are notable taxonomic differences that may occur dependent on the 16S hypervariable region sequenced, however ML models built for ASD classification are possible when utilizing either the V4 or V3-V4 hypervariable regions.

We noted a split in our dataset in regards to sequencing depth, where 6 studies were sequenced at a relatively lower sequencing depth (around 6,000 reads/sample) and 8 were sequenced at a higher sequencing depth (a minimum of 14,000 reads/sample). There was a notable difference in the performance of the model trained on samples from cohorts with a minimum of 14,000 reads per sample, to that of the cohorts with a minimum of 6,000, as assessed by the AUC, Accuracy, and F1 statistic (Fig. 5e). Models trained on data from studies using a high sequencing depth had an average AUC of 0.75 while studies with a low sequencing depth had an average AUC of 0.59. However, from a taxonomic perspective, we found no significant effect of sequencing depth on the relative abundance differences of ASD to controls (Supplementary Figs. 1-4). These results suggest that increased sequencing depth is important for accurate ASD classification, while not impacting taxonomic results.

We next evaluated how aspects related to the demographics of the study participants may alter ASD microbiome results. For the effect of geographic location, we split our cohorts by which country each study’s subjects were from. We found no statistically significant differences in ASD to control relative abundance ratios from studies in the USA vs studies from China (Supplementary Figs. 5-8). However, we did find a striking difference in the ability of GBC models to classify ASD and control samples dependent on country (Fig. 5f). We found our models performed better on Chinese samples with an average AUC of 0.78 vs an AUC of 0.6 on samples from the USA. Our results suggest that geographic location is also an important factor for ASD classification utilizing the gut microbiome.

Some studies utilized age and sex-matched controls, while other studies used household controls such as siblings or mothers. We found no statistically significant differences in ASD to control relative abundance ratios from those with related individuals as controls to those with unrelated individuals (Supplementary Figs. 9-12). We did however observe a notable difference in the GBC model’s ability to classify ASD and controls from the cohorts which utilized unrelated controls to those with related controls (Fig. 5f). Our model performed better from samples of cohorts with unrelated controls with an average AUC of 0.71 compared to cohorts with related controls which had an average AUC of 0.54. This indicates that there are larger differences in the microbial profile of ASD individuals compared to controls when the controls are not related to the ASD individual.

We additionally assessed how sex influenced microbial profiling and the diagnostic accuracy of our ML model of ASD and control individuals. We split our samples by sex and compared the AUC of our model for predicting ASD status of females compared to males (Fig. 5f). We observed a greater accuracy for predicting ASD status from male samples with an average AUC of 0.67 vs an AUC of 0.55 from female samples (Fig. 5f). We did not look at taxonomic differences of ASD vs controls on a per-study basis due to the limited number of females within each dataset. However, the difference in predictive accuracy of ASD status from male samples compared to that of females may indicate the influence of sex on microbial profiling, and subsequent predictive accuracy.

Furthermore, the microbial composition of the gut microbiome is known to shift during human development^49^. We thus assessed how the age of our subjects influenced the ability to predict ASD status. To do this, we subset our data at every potential age range between the ages of 1 and 16 and compared the performance of GBC models in predicting ASD status (Fig. 5g). We observed that samples from the age group of 2-7 years old obtained the greatest predictive accuracy (AUC = 0.69) and that as the age of the individuals increased, the AUC of our model concurrently decreased (Fig. 5g). To validate that our findings were not simply due to the corresponding sample sizes available at each age range, we first determined the effect of sample size on GBC model performance (Extended Data Fig. 7a), then plotted the corresponding sample sizes (Extended Data Fig. 7b). We observed that samples from younger subjects performed better than expected by sample size at ASD status prediction.

With the understanding that study factors strongly influenced our ML performance, we next adjusted our data according to the variability associated with various combinations of study factors and evaluated whether this improved the ML prediction of ASD status. We included variable region, control relationship, country, sex, sequencing depth, and study as “batches” and batch corrected the raw sequence count data for each factor individually, and for all possible combinations of factors. Most combinations of factors significantly increased our baseline GBC model’s performance (baseline AUC = 0.623 ± 0.003) (Fig. 5h). The only batch corrections that did not improve the AUC from baseline were variable region only, control relationship only, or both variable region and control relationship only (Fig. 5h, Extended Data Figure 7c). We suspect adjusting the data by these variables did not improve ML performance due to non-overlapping subsets of features which our batch correction method would not account for. The best performing model contained data batch corrected for variable region, sex, control relationship, and sequencing depth (GBC AUC = 0.687 ± 0.003) (Extended Data Figure 7c). However, what was most notable, was the effect of including sex as a variable in the batch correction (Fig. 5h). The inclusion of sex alone increased the GBC model’s AUC by 2.3% from baseline. Further, the ML performance improved at a faster rate when more variables were included alongside sex (Fig. 5h). While our best performing model does not reach an AUC comparable to state-of-the-art ASD classifiers built from brain imaging data (AUC 0.74)^50^ or behavioral videos (AUC 0.83)^51^, these results highlight a strong connection between ASD and the microbiome, as well as how study factors can influence the strength of this connection.

## Discussion

In the current study we employed systematic approaches to interrogate the existing ASD gut microbiome data and provide an in-depth report of the commonalities and discrepancies between datasets. We further tested the extent to which study design attributes influence microbiome associations to ASD as well as the accuracy of machine learning models designed for ASD classification and prediction. Given the vast variety of choices in different data processing approaches, and the variability that each of these choices introduces^52^, we utilized a consistent methodology to minimize technical effects between studies. With our dataset of over 1,700 samples, we addressed the question of whether, and how, the ASD microbiome differs from control individuals. We further investigated new questions in the field by analyzing the data based on study-specific factors including; 16S hypervariable region, control group demographics, age, sex, and country of origin. Importantly, we also utilized eleven different ML models to evaluate the impact each of the above factors has on the classification and prediction of ASD status using microbiome data. Our results have thus provided a reference point for the ASD microbiome field while highlighting important considerations for future ASD related studies and their experimental designs.

Our study first addressed commonly reported metrics within the microbiome field by evaluating the ASD microbiome for differences in alpha-diversity, beta-diversity, and changes in the relative abundance of the microbiome at different taxonomic levels. We observed relatively little consistency in identifying alpha-diversity differences, though half of the studies found a significant difference in at least one of the four metrics tested. Significant beta-diversity differences between ASD individuals and controls were relatively more common and evident within the total dataset. These results could indicate that there is a distinct microbiome within the ASD population, but the diversity within their microbiome is not a distinguishing factor. Therefore, the alpha-diversity differences observed within some studies may be the result of additional variables outside of ASD status.

Regarding what microbes may be driving the differences between ASD and controls, large study variability was a hinderance. Despite this, we were able to identify several consistent trends in the ASD gut microbiome across cohorts. The largest taxonomic consistency was the enrichment of Actinobacteria, at both the phylum and class level as well as increased *Bifidobacterium*, and a general decrease in *Prevotella* among ASD individuals. The genus *Bifidobacterium* (phylum Actinobacteria) are important to human development early in life, while they are later replaced by strict anaerobic bacteria^53^. It is possible that the increased prevalence of *Bifidobacterium* is indicative of an underdeveloped gut microbiome. Interestingly, the idea that the ASD gut microbiome is underdeveloped was supported by a recent metagenomic study of ASD, where they found gut developmental associated microbes absent in ASD individuals compared to age matched controls^54^.

Microbial compositions are known to vary across different human populations^27^. Additionally, the various techniques used to analyze the microbiome can impact the estimated biological diversity^52; 55^. Our study contextualizes the ASD microbiome field by evaluating how both technical choices and cohort demographics alter reported results. Of interest, we observed significant discrepancy in the diagnostic capability of our ML models based on data from geographic regions, i.e. the USA and China, and observed increased predication rates in the samples collected from the latter country. The underlying reasons are unknown, but several explanations may exist. Firstly, the ASD samples from our Chinese cohorts may represent more severe cases of ASD than samples from the western studies. Alternatively, our results may also be a result of different countries representing populations that differ in socio-economic status, diets, living conditions, and environmental factors. Unfortunately, we were unable to obtain sufficient clinical and diagnostic metadata from our cohorts, and were thus unable to directly test whether ASD individuals from China represent more severe cases than those from the USA.

Of particular interest, we observed significantly increased accuracy of our ML algorithms to predict ASD status within an early age window (2-7 years). This increased accuracy has several implications for both the development of microbiome-based diagnostics, but also the importance of age in assessing the microbiomes’ role in the pathogenesis of ASD. Currently, diagnosis of ASD is often delayed partly because standardized diagnosis involves the identification of abnormal behaviors that may emerge after the disease is established^56^. Thus, development of early detection diagnostics is severely needed. The capacity of ML algorithms to detect ASD status from a young age highlights the potential of the microbiome to facilitate early diagnoses. Further, our findings of increased accuracy from younger children suggests a starting age that is consistent with the establishment and maturation of the enteric microbiota^57^, potentially indicating a critical window of the microbiome in the pathogenesis of ASD. However, many confounders exist, including bias in the studies that samples belonged to or the potential that ASD cases at ages 2-7 represent more severe cases than those of older children.

The bacterial 16S rRNA gene spans the hypervariable regions V1-V9. Depending on the particular amplification primer sets that are used, researchers can target any particular region. While some large-scale efforts such as the earth microbiome project endorse the amplification of the V4 region, a standard in the field is lacking^58^. This lack of standardization has been shown to impact reproducibility especially in metrics estimating biological diversity^52; 55^. We found that the 16S hypervariable region of choice indeed contributed to cohort discrepancies regarding taxonomic associations to ASD. We observed discordant results in both per-study and per-sample relative abundances for the ratios of the genera *Prevotella* to *Bacteroides*, genus *Prevotella*, order *Desulfovibrionales*, and class *Deltaproteobacteria* in V3-V4 versus V4-only cohorts. It was particularly interesting that *Prevotella* appeared to be increased in ASD among V4 datasets while it was decreased in ASD among V3-V4 cohorts. The V3-V4 region represents a larger portion of DNA (∼400 bp vs ∼150 bp among these studies). A potential explanation could exist in the varying taxonomic assignments of reads based on longer or shorter read lengths. Therefore, our results indicate that a portion of the study discrepancies in the field may be caused by differences in 16S sequencing methods.

Other factors our study identified as leading to discrepancies among ASD microbiome cohorts include sampling depth and control type. While we did not note a significant impact on the taxonomic outcomes due to these study design choices, we observed significantly improved ML performance among studies which performed deeper sequencing and utilized unrelated controls. These findings suggest that ASD households may have distinct microbiomes from the general population, and this factor should be accounted for within future studies. Our sequencing depth results suggest that deeper sequencing may be important for future diagnostic development based on the ASD microbiome.

In addition, we show that by accounting for microbiome variability related to study parameters, we were able to significantly improve ML prediction of ASD status. Interestingly, the variable which increased ML accuracy upon normalization the most was accounting for differences in sex. Underlying this analysis, our batch correction method adjusted the mean and standard deviations of male and female samples, suggesting that there exists sex-specific variability in microbiome compositions which is relevant to ASD status.

Overall, our best performing ML model was based on Chinese samples (AUC = 0.78) achieving performance better than state-of-the-art models built from brain imaging data (AUC = 0.74)^50^ but not behavioral videos (AUC = 0.83)^51^. When including all samples, our best model was based on data which removed microbiome variability related to variable region, sex, control relationship, and sequencing depth. This model had strong performance at ASD classification, though the performance was worse than state-of-the-art ASD classifiers (AUC = 0.69). These findings suggest that ASD diagnosis based on the microbiome may be more accurate for some sub-populations than others.

Regardless of the study discrepancies described above, we were particularly interested in a consistent theme we observed among the predicted functional discrepancies found within the ASD gut microbiome. Chorismate is a common branch point for the synthesis of several biologically relevant molecules including the AAA’s, menaquinones, ubiquinones, enterobactin, and folate. We found that many of the pathways commonly different within the ASD microbiome were related to metabolites downstream of chorismate. Intriguingly, many of these molecules have been discussed individually, but not in concert in ASD-related literature. For example, folate levels are associated with ASD^59^ and folinic acid treatment has shown promise for treating language impairment in ASD^60^. Further, tryptophan levels (an AAA) are frequently reported to be dysregulated in ASD^10; 54^, and could be related to ASD pathology through downstream metabolites, serotonin and kynurenines^61^. Further, the breakdown of tyrosine (another AAA) to 4-ethylphenyl sulfate (a molecule increased among ASD subjects) was recently shown to be linked to anxiety in mice^62^. Intriguingly, chorismate biosynthesis is dependent on the enzyme 5-enolpyruvylshikimate-3-phosphate synthase (EPSPS), which is the target of glyphosate. Although a debate still exists, some researchers have linked glyphosate levels to ASD rates and an altered microbiome^63^. Our results suggest that ASD could be associated with, or a result of, a disturbance in the gut microbiota’s role in producing or modulating various downstream, bioactive products of chorismate. While this data was of high interest, it is important to note that we observed limited significance in these differences as well as variability between studies. Moreover, these results are based on predicted functions, and need to be validated by multi-omics approaches including shotgun metagenomics and metabolomics.

Finally, there are several limitations in our work. First, we were limited to publicly available data which rarely included adequate clinical data to evaluate alternative hypotheses. Absent of this information, we could not determine whether the inconsistency between studies simply reflects a sampling bias of selected studies. For instance, some studies may only enroll individuals with more severe forms of ASD whose microbial profiles may be distinct from those of neurotypical individuals. Further, some studies enrolled subjects currently experiencing varying states of gastrointestinal distress such as constipation, a common comorbidity of ASD^20^. Studies with these types of sampling biases may artificially inflate the importance of various metadata variables (e.g. country, variable region, sequencing depth). Thus, although we speculate that the observed metadata variables drive the differences in predictability of ASD between studies, an alternative hypothesis is that unobserved metadata variables (e.g. phenotypic severity) are driving these differences. Second, our methods often compared microbial compositions. While we found this to be an important baseline analysis, accounting for compositionality through either computational or laboratory controls might reveal more consistent associations between ASD and control subjects^64^. Third, our study utilized only 16S data, which is limited in its ability to reliably resolve taxonomy beyond the genus level. Further studies with multi-omics approaches are needed to quantify microbial species abundances, and the proteins and metabolites they produce.

In total, our current study represents an important synthesis of the state of the ASD microbiome field. We report novel factors that likely explain the variabilities existing in the field, including the age and country of study participants as well as the variable region targeted and choice of control subjects. We observed significant differences overall between ASD and control subjects, though notable variation exists between studies. While our study includes over 1700 subjects, the observed variability in ASD due to the factors identified by this study warrants further investigation from large cohorts that span multiple geographic locations, age ranges and control types.

## Methods

### Literature search strategy and selection criteria

We performed a systematic literature search through the PubMed database, from inception to July 15, 2021, to identify applicable studies assessing autism spectrum disorder and the gut microbiome. The following search terms were used as free-text words only, or as key words; ASD, or (Autism Spectrum Disorder, or Autism), and (gut microbiome, or microbiota, fecal microbiome). No other search features, or advanced language limits were used. Articles were included in this meta-analysis if; (a) fecal microbiota samples were taken of a patient with autism and or that of a typically developing individual (b) data and respective metadata were retrievable and interpretable (c) autism diagnosis was given by a healthcare professional. From our literature search, a total of 244 studies were assessed for inclusion. Of the 244, 36 studies indicated cohorts wherein the microbiome’s role in ASD was researched and were read in full to assess their inclusion eligibility. A total of 12 met the inclusion criteria described above and were included in this analysis. We identified one dataset deposited in the Sequence Read Archive (SRA) with its respective metadata, that was not yet published but published during manuscript preparation^39^. Lastly, we utilized open-source data generated from the American Gut Project (AGP) to cultivate an additional dataset of ASD children and controls. We designed a propensity score algorithm (see below) to identify a best-match control subject for each ASD subject. Together, these 14 cohorts yielded a total of 888 ASD samples and 852 control samples of typically developing individuals from a wide range of ages, and geographic locations.

### Exclusion criteria

With our goal to minimize known technical effects we did not include studies which only utilized shotgun metagenomic sequencing methods, Roche 454 sequencing, and or 16S rRNA pyrosequencing. Thus, 15 studies were excluded from our analysis for incompatible sequencing methods. Eight other studies met our inclusion criteria, but did not have publicly available metadata. All eight authors were contacted but did not respond to our request for data access and thus were unable to be included in our analyses.

### Compiling a cohort from the American gut project

The AGP dataset contains samples from >29,000 participants thus representing a diverse dataset to compile targeted cohorts with. All available 16S data and metadata was downloaded from Qiita.ucsd.edu (https://qiita.ucsd.edu/study/description/10317). As these samples were part of a citizen-science initiative where samples may have been exposed to variable collection conditions, we first applied a bloom filter (https://github.com/knightlab-analyses/bloom-analyses) to remove Gammaproteobacteria species known to commonly grow at room temperature^65^.

Within this dataset, we identified fecal samples from subjects which were reported to be diagnosed with ASD by a physician and next compiled a control cohort based on each ASD subject’s features. Therefore, we adapted a method based on propensity score matching which consists of two main steps: scoring and matching^66^. For each sample in the ASD cohort, a corresponding control sample was selected based on having an exact match of specific metadata features (run date, sample type, and sex) and having the closest “similarity score” calculated from other metadata features (age, BMI, c-section, country, diabetes, antibiotic history, probiotic frequency, prepared meals frequency, allergies, prepared foods frequency, red meat frequency, fermented food consumption, whole grain frequency, vitamin B supplementation frequency, plant protein frequency, vitamin D frequency, vegetable frequency, epilepsy or seizure disorder). Exact matching allows us to select features that need to be controlled for. For example, to control for technical variability between sequencing runs, we required the control sample to have come from the same sequencing run date as the ASD sample. On the other hand, the similarity scores are simply the conditional probability of cohort assignment given a vector of metadata features. In this way, the matched control samples represent a subset of the control cohort that have similar, if not the same, metadata feature distributions as that of the ASD samples.

To calculate similarity scores, each feature was first binarized using the OneHotEncoder class from scikit-learn version 0.24.2 (https://scikit-learn.org/). These features were then fit using scikit-learn’s LogisticRegression class where the dependent variable was cohort status (i.e. 1 for ASD cohort; 0 for control cohort). The similarity score is therefore simply the output of the sigmoid function which is also the predicted probability of being assigned to the ASD cohort.

During the matching process, the similarity score from each sample in the ASD cohort is then compared with the similarity score from every control sample whose “exact matches” variables were the same. The control sample with the closest similarity score is then placed in the control cohort and removed as a possible control sample for subsequent rounds of matching. After matching every ASD sample, we obtain a control cohort of unique samples and whose sample size is the same as that of the ASD cohort serving as our ASD and control samples for the AGP dataset.

### Data Acquisition

For data publicly deposited in the NCBI Sequence Read Archive (SRA), sample metadata was downloaded from each project using the SRA Run Selector tool (https://www.ncbi.nlm.nih.gov/Traces). A custom script (process_experiment.py) available on github (https://github.com/mortonjt/GetData) was used to systematically download, trim primers and process samples into data tables. In cases when data and sample metadata was not deposited through SRA, it was provided directly from the original study’s authors and processed using identical parameters as described below.

### Standardized 16S processing pipeline

Our goal was to collect and reprocess each study in a systematic and harmonious way to first, reduce errors imposed by different processing methods, and second to enable us to better draw conclusions about the ASD gut microbiome and our dataset as a whole. We therefore processed each study individually utilizing the same processing steps with QIIME 2 (v. 2020.8)^67^. We additionally processed all the samples together with the same pipeline, to draw conclusions and uncover trends about the aggregated data as a whole. The first step of our systematic processing pipeline was the removal of primers when needed utilizing the custom script process_experiment.py. Next, all sequences were trimmed to a length of 150 bp through Deblur^68^. After trimming, sequences were denoised and filtered using the Deblur QIIME 2 2020.8 plugin^68^. Taxonomic classification was performed with the QIIME 2 q2-feature-classifier plug-in with the full length 16S pre-fitted GreenGenes-trained Naive Bayes classifier (gg-13-8-99-nb-classifier.qza) provided by QIIME 2 (QIIME 2 v. 2020.8)^69; 70^. Phylogenetic placement was assigned with the SEPP fragment insertion plug-in from QIIME 2 (v. 2020.8)^71-73^. SEPP was chosen to minimize the effects of hypervariable regions by assigning short reads to a reference phylogeny^71^. All features that were not aligned to the tree were removed with the q2-fragment-insetion filter-features QIIME 2 2020.8 plugin^71^. The resulting tables were ASV-level read counts which were used for all downstream analyses.

Prior to alpha and beta diversity analyses, the sequences from each cohort were rarefied independently to the necessary depth to retain the maximum number of reads as possible, while filtering any samples where the number of reads was very low (see the available code for exact parameters and respective sequencing depths). For alpha and beta-diversity analyses of the aggregated data, the resulting table was rarefied to a sequencing depth of 6000 with the core-phylogenetic QIIME 2 v. 2020.8 plug-in. Alpha diversity was assessed with Faith’s phylogenetic diversity, Shannon diversity, species evenness, and observed features from the core-phylogenetic QIIME 2 v. 2020.8 plug-in^74^. The unweighted UniFrac and weighted UniFrac distance matrices, also calculated using the core-phylogenetic QIIME 2 v. 2020.8 plug-in, were used to compute beta diversity distances and corresponding Principle Coordinates Analysis (PCoA) plots^41; 75^.

### Analysis of alpha-diversity, beta-diversity, taxonomic and functional differences

We filtered all samples that contained less than 6,000 reads per sample before all fold change, relative abundance, pathways abundance, and ML comparative analyses, for both the aggregated data and per-study level analysis. Analyses were performed using custom Python (version 3.6.0) scripts. Statistical significance of beta-diversity distances between groups was calculated using the QIIME 2 beta-group-significance and reported p-values were calculated using Permutational multivariate analysis of variance (PERMANOVA). Differences between ASD and control subject’s alpha-diversity was performed only on a per-cohort basis due to large variability between studies. The alpha diversity log2 ratio of ASD to controls was performed in NumPy version 1.19.1 (https://numpy.org/), from ASD and control subjects mean values as calculated using SciPy version 1.5.2 (https://scipy.org/). The significance of the log2 fold change for each cohort was assessed by an unadjusted, two-tailed t-tests with assumed unequal variance in SciPy version 1.5.2.

For the per-study log2 fold changes of each taxonomic level mentioned, samples were first normalized to relative abundances by dividing ASV read counts by the total reads per sample. Relative abundances were summed at each taxonomic level using Pandas version 0.25.3 (https://pandas.pydata.org/). At the per-study level, log2 ratios of ASD and control subject means were calculated using SciPy version 1.5.2, and log transformed in NumPy version 1.19.1. Statistical significance was assessed in the same manner as that for alpha diversity described above. The log10 transformed relative abundance in each sample for the significant taxa we highlighted was performed in Pandas version 0.25.3. To avoid instances of infinity due to log10 transformation, the minimum non-zero normalized relative abundance value was added to all samples utilizing NumPy version 1.19.1. Next, we log10 transformed each samples normalized relative abundance with NumPy version 1.19.1 before plotting.

Microbial functional profiles were estimated from 16S data using the Phylogenetic Investigation of Communities by Reconstruction of Unobserved States (Picrust2) software (v. 2019.1)^76^. The log2 fold change of ASD/controls of the resulting pathway abundance tables were calculated in python as described above. The geom_point plot of the log2 fold change values were plotted with ggplot2 version 3.3.3 in R version 4.0.5. A clustermap using the seaborn package version 0.11.0 was generated and used to reorder the geom_point plot in R. The dendrograms from the seaborn clustermap and applied to the geom_point plot in R when present. All figures were generated using ggplot2 version 3.3.3 in R version 4.0.5. A full list of the R packages used, along with their versions can be found in Extended Data Table 3.

### Machine learning classification of ASD status

To predict ASD status from the microbiome, we used eleven different classification models, evaluation metrics, and auxiliary functions from the scikit-learn package version 0.24.2 (https://scikit-learn.org/). Unless stated otherwise, we used 5-fold cross validation repeated 10 times to estimate the averages and variances of different performance metrics. For a complete list of models and their corresponding hyperparameters, see Extended Data Table 2. We did not perform hyperparameter optimization. Reported results include the area under the curve (AUC) of the ROC curve, as well as the accuracy and the F1 score of each model.

During our initial evaluation of different ML models, we used the aggregate dataset summing reads by the highest taxonomic annotation given to each ASV. The dataset was normalized so that relative abundances sum to unity before training and evaluating each ML model. Next, for each study (including the aggregate dataset), we collapsed the dataset at different levels of taxonomical hierarchy (e.g. genus) by grouping each taxa (e.g. *Clostridium*), summing their raw read counts, and normalizing so that the relative abundances sum to unity. This processed dataset was then used for training and testing the GBC, ABC, and the decision tree classifier from scikit-learn. Reported results include the area under the curve (AUC) of the ROC curve, as well as the accuracy and the F1 score of each model.

To determine the importance of each feature, we applied the Shapley additive explanations (SHAP) package (https://github.com/slundberg/shap) to the Gradient Boosting Classifier model trained on the aggregate dataset^77^. The top 25 most important features by cumulative SHAP value were displayed using the beeswarm plotting function of the SHAP package. Absolute SHAP values from each of the top 25 features were averaged for each study and displayed as a clustermap using the seaborn package version 0.11.0 (https://seaborn.pydata.org/). To determine the effects of different metadata variables, we separated the entire dataset by their corresponding metadata features. Each partition of the dataset was further split using 5-fold cross validation and this data was used to train and evaluate a Gradient Boosting Classifier trained to predict ASD or control status. The AUC of ROC, F1 score, and accuracy were used to evaluate model performance. The entire process was repeated 50 times to estimate the standard error of each performance metric.

To estimate the predictability of ASD at different age ranges, we evaluated the predictability of ASD across different age ranges from 1-to 16-year-olds. That is, we used only the subset of data that fell into each age bin (e.g. 1-3 year olds). To ensure sufficient data within each age bin, the lower and upper bounds of each bin had to be separated by 2 years. Using 5-fold cross validation of this subset of data, we trained and evaluated 30 instances of GBC models. The average AUC of the ROC curve of each age bin was displayed using a seaborn heatmap. Age ranges that were invalid (e.g. 3-1 year olds) were excluded and displayed as a gray square.

To explore how correcting for different metadata variables in our dataset can improve model performance, we used pycombat version 0.20 (https://github.com/CoAxLab/pycombat) – a python implementation of the Combat algorithm^78^, to perform batch correction. We included “16s variable region”, “control relationship”, “country”, “sex”, “sequencing depth”, and “study” as metadata variables to account for during batch correction of sequencing count data. In addition to considering each of these variables individually, we also explored the possibility of an interaction effect between these variables by evaluating every subset of the power set of these variables. For every subset in consideration, we first performed one-hot encoding of every unique combination of metadata variable values in the subset. Then, we performed batch correction using pycombat, and then, split the data into training and testing sets. Lastly, we trained a GradientBoostingClassifier on the training set and evaluated the area under the ROC curve (AUC) of the model’s predictions of the testing set. We performed this entire process 50 times to generate confidence intervals.

## Data availability

Data utilized in this study was from publicly available datasets deposited in the Sequencing Run Archive (SRA, https://www.ncbi.nlm.nih.gov/sra). Data can be accessed using the following study identifiers: PRJNA282013, PRJNA578223, PRJNA642975, PRJNA453621, PRJNA644763, PRJNA687773, PRJNA624252, PRJEB27306, PRJEB27306, PRJNA529598, PRJEB42687. Other datasets included data from David et al. 2021 available at http://files.cgrb.oregonstate.edu/David_Lab/ASD_study1/ and data from the AGP available from Qiita under study identifier 10317 (https://qiita.ucsd.edu/study/description/10317).

## Code availability

Data processing scripts are available on Github (https://github.com/mortonjt/GetData). Notebooks for both data analysis, and visualizations are also available on Github (https://github.com/precidiag-inc/ASD_16S_metaanalysis).

## Acknowledgements

We thank James Morton and Gaspar Taroncher of the Simons Foundation for assisting in data collection and processing, helpful conversations surrounding the project and for reviewing the manuscript. We further acknowledge the R&D members of Precidiag Inc., including Zhuoteng Yu, Rhys Duqeutte and Hang Yin, for helpful discussions and the administrative team members for research support. We thank the Flatiron Institute and the Commonwealth Computational Cloud for Data Driven Biology for donating supercomputing resources toward this project. We also thank Professor Dong Kong (BCH/HMS) for their advice and comments regarding the manuscript. We also thank Chris McDougle and Robyn Thom (MGH/HMS) for comments regarding the project.

## Contributions

R.H.M. conceived and designed the study. A.C., R.H.M., E.H.W., developed the methodology. R.H.M. acquired the data. A.C., E.H.W., R.H.M. analysed the data. A.C., E.H.W., R.H.M. interpreted the data. A.C., E.H.W., R.H.M. wrote the manuscript. All authors reviewed and revised the manuscript.

