## Supplementary Information for "Meta-analysis of the autism gut microbiome identifies factors influencing study discrepancies and machine learning classification"

**Extended Data Tables**

**Extended Data Table 1. Characteristics of included studies.** Details regarding each included study are compiled including the number of samples analyzed, age ranges with mean and standard deviations, the percentage of males included, country, control type and sequencing details.

**Extended Data Table 2. Hyperparameters used in machine learning analyses.** Table of machine learning models from the scikit-learn package and their corresponding hyperparameters. Default hyperparameters were used unless specified otherwise.

| Package | Model | Hyperparameters |
| --- | --- | --- |
| scikit-learn | K Neighbors | n_neighbors = 3 |
| | Support Vector Classification (SVC Kernel) | kernel = linear<br>C = 0.025 |
| | Support Vector Classification (SVC Gamma) | gamma = 2,<br>C = 1 |
| | Decision Tree | max_depth = 5 |
| | Random Forest | max_depth = 5,<br>n_estimators = 10,<br>max_features = 1 |
| | Multilayered Perceptrons (Neural Networks) | alpha = 1,<br>max_iter = 1000 |
| | Gaussian Naïve Bayes | |
| | AdaBoost | |
| | Gradient Boosting Classifier | |
| | Quadratic Discriminant Analysis | |

**Extended Data Table 3. R Packages and Version Number Utilized For Plotting.** Table of R version 4.0.5 packages and their corresponding version numbers used for plotting all figures.

| Package | Version number |
| --- | --- |
| plyr | 1.8.6 |
| hrbrthemes | 0.8.0 |
| gcookbook | 2.0 |
| gapminder | 0.3.0 |
| tidyverse | 1.3.1 |
| viridis | 0.6.1 (Neural Networks) |

| | |
|---|---|
| ggrepel | 0.9.1 |
| ggplot2 | 3.3.3 |
| forcats | 0.5.1 |
| gridExtra | 2.3 |
| ellipse | 0.4.2 |
| rstatix | 0.7.0 |
| dplyr | 1.0.6 |
| ggpubr | 0.4.0 |
| purrr | 0.3.4 |
| wesanderson | 0.3.6 |
| readr | 1.4.0 |
| reshape2 | 1.4.4 |
| tidyr | 1.1.3 |
| extrafont | 0.17 |
| RColorBrewer | 1.1-2 |
| tibble | 3.1.2 |

### Extended Data Figures

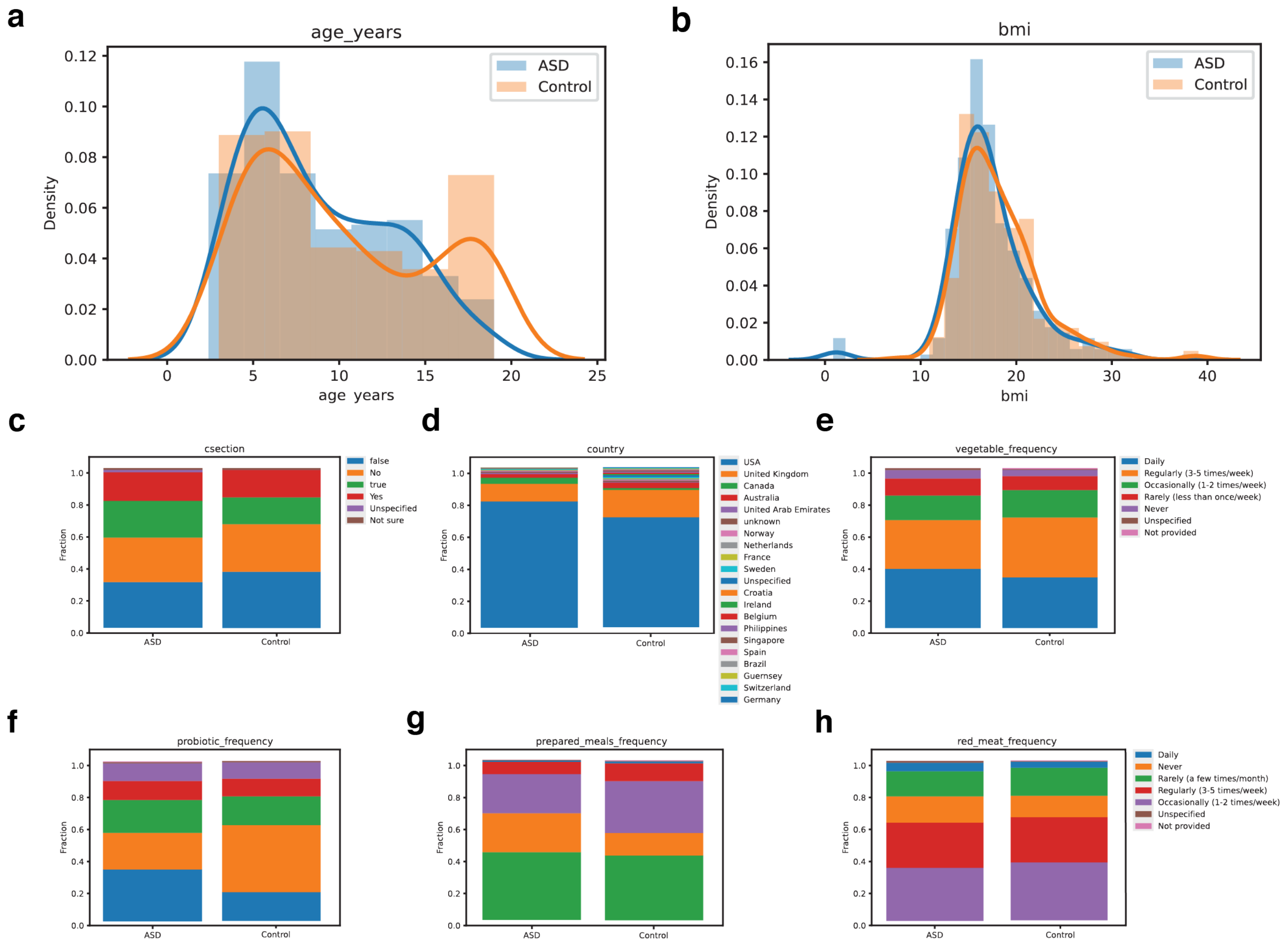


**Extended Data Figure 1. Comparison of ASD and control demographics from data compiled from the American gut project. (a-b)** Distribution of ages (a) and the BMI (b) of ASD and control subjects compiled from the AGP. Kernel density plots are shown overlaying histograms. **(c-h)** Distributions of categorical demographics from ASD and control subjects compiled from the AGP. Stacked barplots showing the fraction of each entry are shown for c-section as a mode of delivery (c), country of origin (d), frequency of vegetable consumption (e), frequency of probiotic consumption (f), frequency of prepared meal consumption (g), and red meat consumption (h).

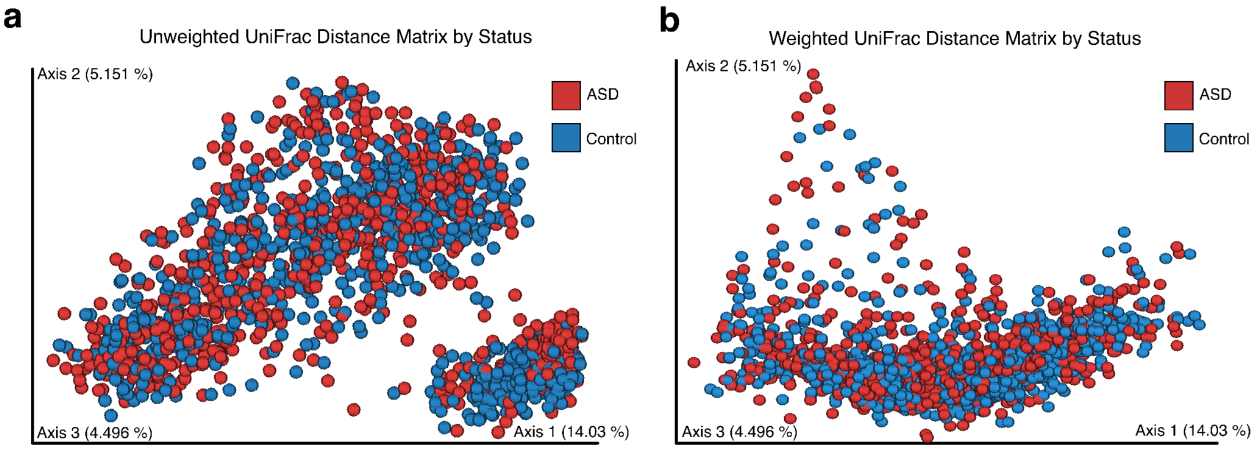


**Extended Data Figure 2. Principal coordinate analysis of samples by status. a,** Principal Coordinate Analysis (PCoA) of the unweighted UniFrac distance matrix of samples rarefied to 6000 reads per sample and colored by ASD status (n = 1492). **b,** PCoA of the weighted UniFrac distance matrix of samples rarefied to 6000 reads per sample and colored by ASD status (n = 1492).

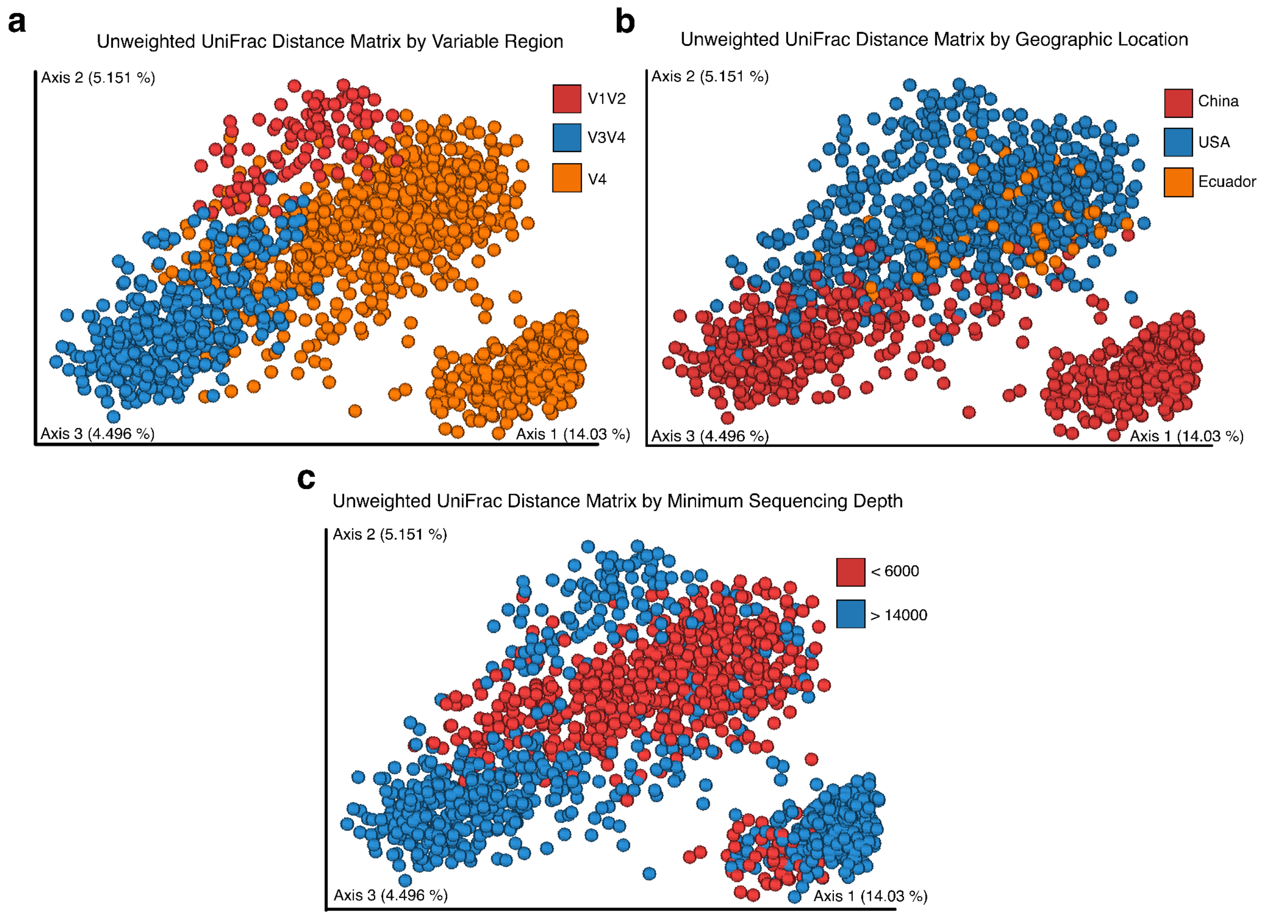


**Extended Data Figure 3. Principal coordinate analysis of samples colored by study design factors. a,** PCoA of the unweighted UniFrac distance matrix of samples rarefied to 6000 reads per sample and colored by hypervariable region (n= 1492). **b,** PCoA of the unweighted UniFrac distance matrix of samples rarefied to 6000 reads per sample and colored by geographic location (n = 1492). **c,** PCoA of the unweighted UniFrac distance matrix of samples rarefied to 6000 reads per sample and colored by sequencing depth (n = 1492).

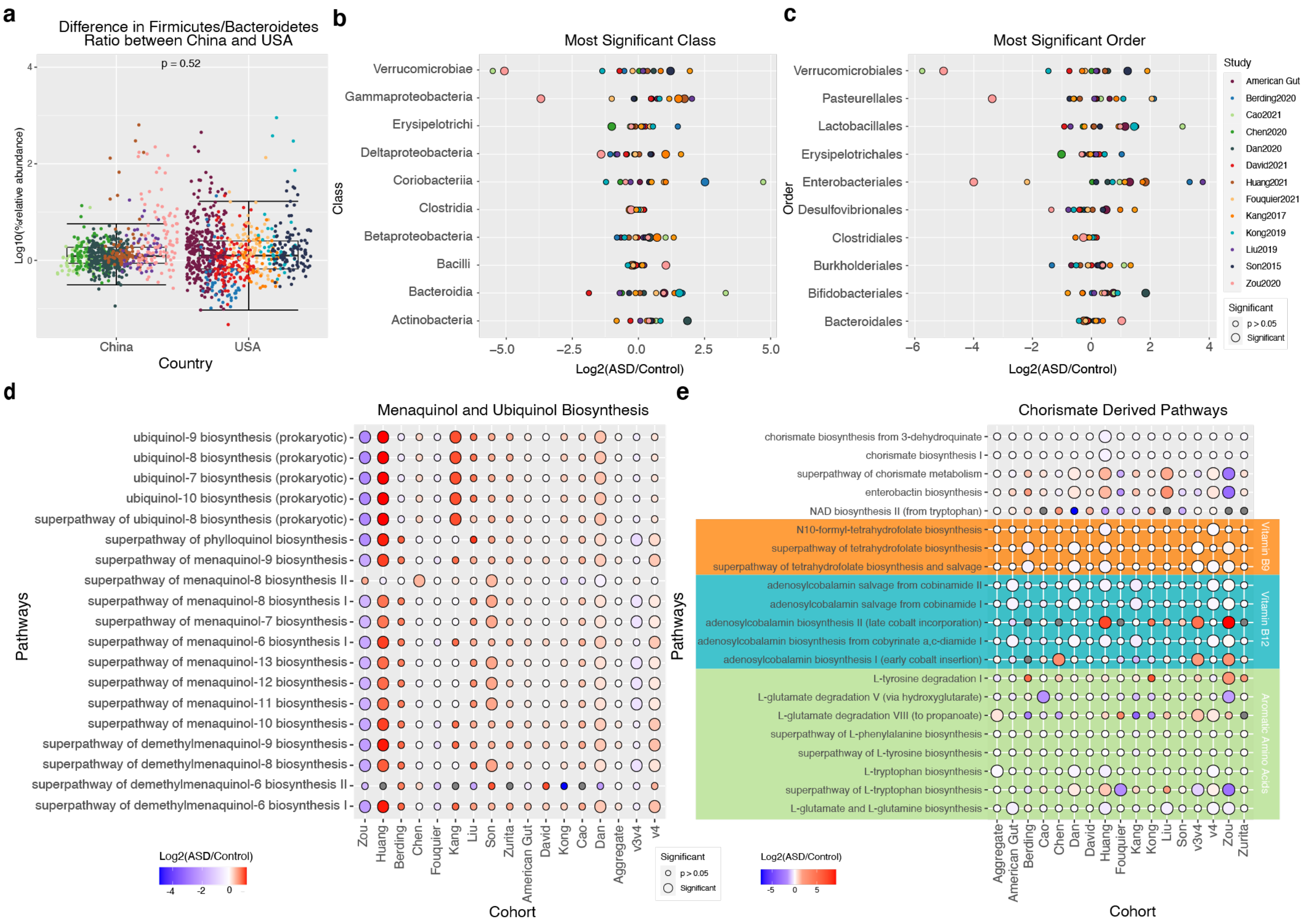


**Extended Data Figure 4. Extended taxonomic and functional differences between ASD and controls by cohort. a,** Log10 transformed relative abundance of each sample stratified by country, and colored by cohort (China n = 678, USA n = 755). A Wilcoxon two-tailed t-test was performed between groups (P = 0.32). Boxplots show the median, quartiles and 1.5x inter-quartile range of the data distribution. **(b-c)** Per-study Log2 fold change in the relative abundance of ASD/controls of the most commonly significant bacteria at the class (b) and order (c) level for each cohort processed individually. Individual point size indicates statistical significance between ASD and controls (unadjusted two-tailed t-tests of unequal variance p-value < 0.05). **d,** Normalized log2 fold change values of ASD/control for every pathway involving menaquinones and ubiquinones in each cohort processed individually or in aggregate. Individual points are colored by Log2 ASD/control fold-change and sized to indicate statistical significance between ASD and controls (unadjusted two-tailed t-tests of unequal variance p-value < 0.05). **e,** Normalized log2 fold change values of ASD/control for chorismate, aromatic amino acids, and vitamin B pathways from each cohort processed individually or in aggregate. Individual points are colored by Log2 ASD/control fold-change and sized to indicate statistical significance between ASD and controls (unadjusted two-tailed t-tests of unequal variance p-value < 0.05). Pathways outlined in green are related to aromatic amino acids, and pathways outlined in blue are vitamin B related pathways.

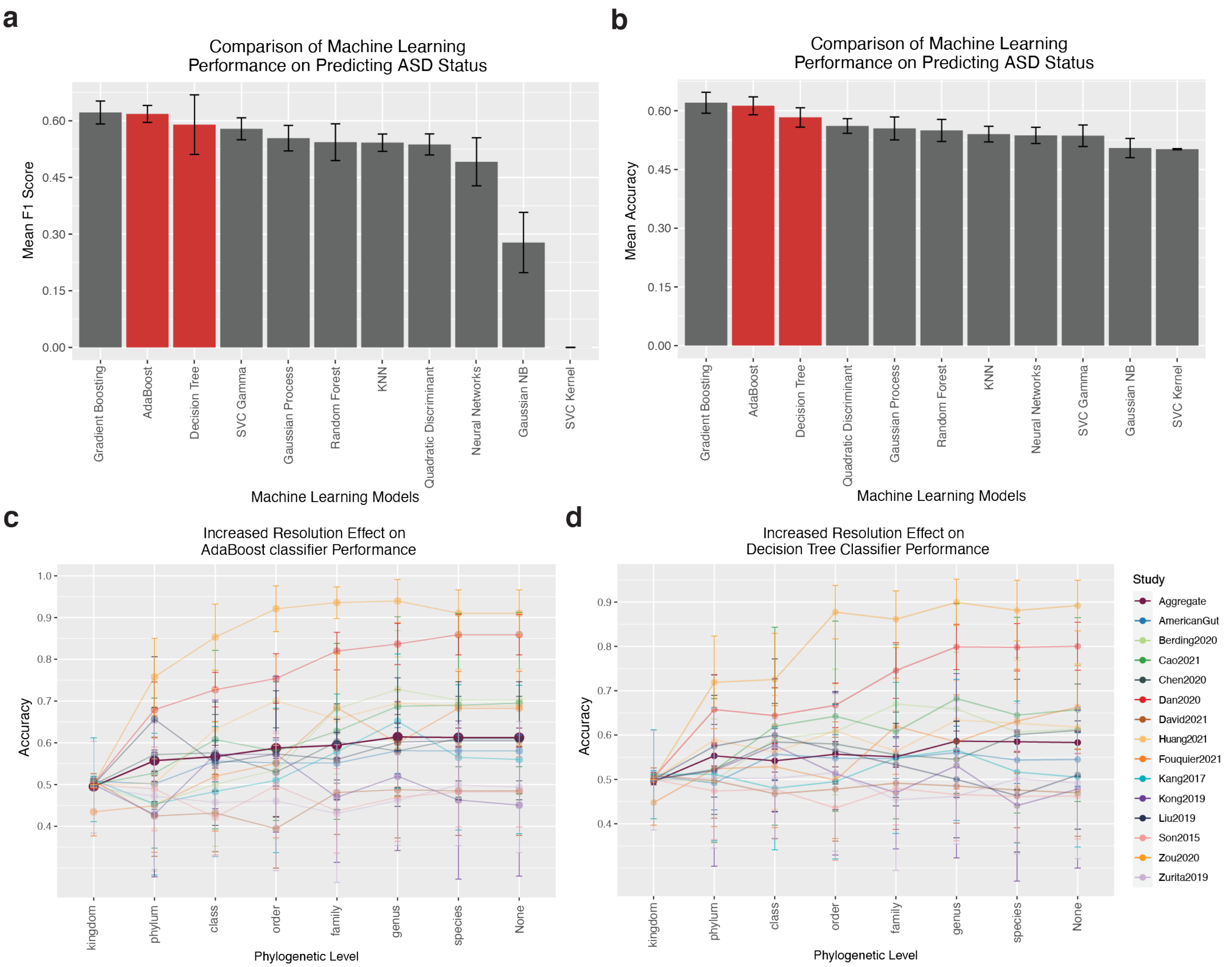


**Extended Data Figure 5. Adaboost and decision tree classifier performance on ASD status prediction.** **(a-b)** Barplots of the average F1 statistics (c) and accuracy (d) +/- SEM, from five iterations of train test-splits in decreasing order, on the aggregated data of ASV's given the most resolved taxonomic assignment for eleven machine learning classifiers. Highlighted in red are the 2$^{nd}$ and 3$^{rd}$ best performing models which are further tested in (a-b). **(c-d)** The mean accuracy of the AdaBoost classifier (a) and Decision Tree classifier (b) models trained and tested on both the aggregated data, and each study individually on reads counts collapsed at different taxonomic levels. Error bars generated from 5-fold cross validation repeated 10 times at each taxonomic level.

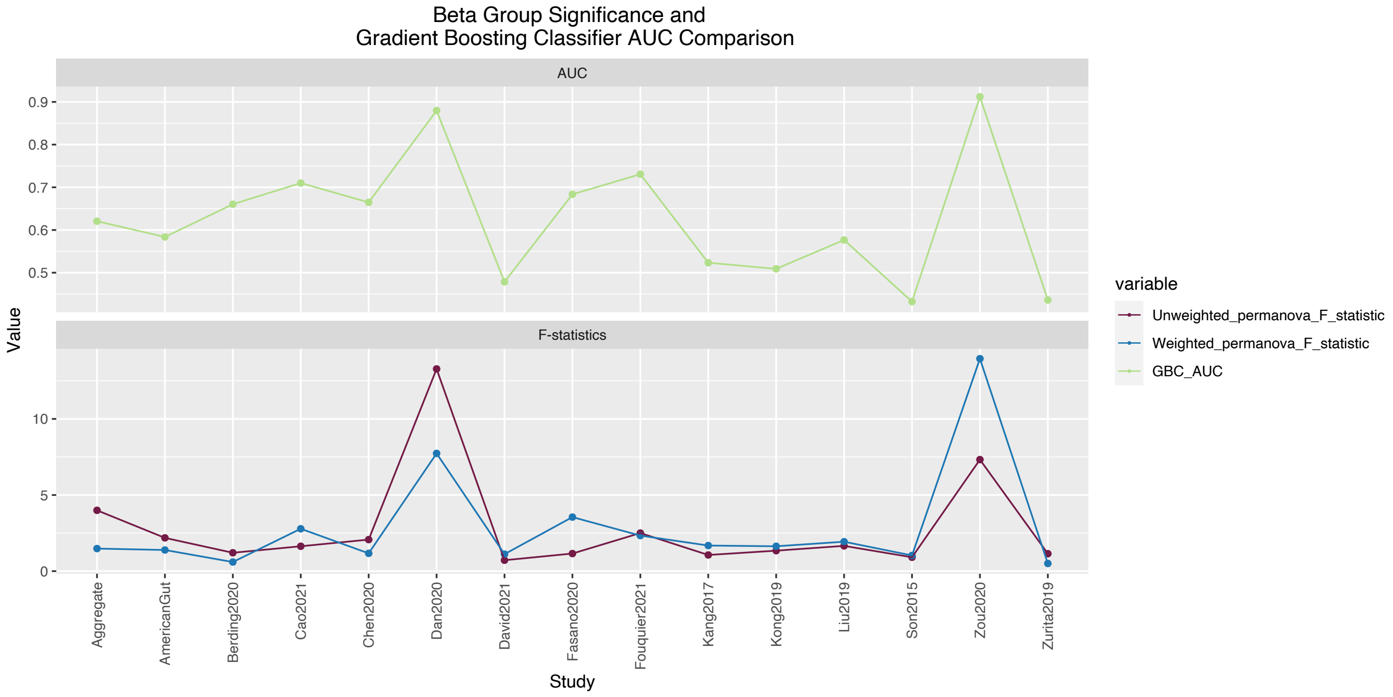


**Extended Data Figure 6. Comparison of beta-diversity results and machine learning results.** Line plot of the unweighted and weighted Pseudo-F statistic for each cohort's beta group-significance test for ASD status, along with the AUC for the Gradient Boosting Classifier (GBC) AUC. The Pseudo-F statistic value was calculated from a PERMANOVA beta-group significance test of ASD vs control from unweighted and weighted UniFrac distance metrics. The GBC model was trained and tested on each cohort individually on reads counts from ASV's at the most resolved taxonomic assignment, and the AUC's are reported. The unweighted and weighted UniFrac Pseudo-F statistics were both positively correlated with the GBC AUC trained on each cohort independent (Pearson correlation coefficient; weighted UniFrac and GBC AUC = 0.79, unweighted UniFrac and GBC AUC = 0.75).

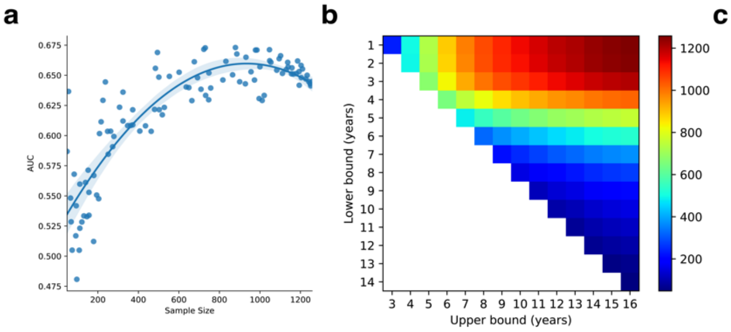


c

| Technical Effects Included in Batch Correction | AUC ± SEM |
|---|---|
| Variable_Region, Sex, Control_relation, seq_depth_range | 0.687 ± 0.003 |
| Country, Sex, Study | 0.685 ± 0.003 |
| Sex, Control_relation, Study | 0.685 ± 0.003 |
| Variable_Region, Country, Sex, seq_depth_range, Study | 0.684 ± 0.003 |
| ***Aggregate (no batch correction)*** | ***0.632 ± 0.003*** |
| Variable_Region, Control_relation | 0.630 ± 0.003 |
| Variable_Region | 0.625 ± 0.003 |
| Control_relation | 0.623 ± 0.003 |

**Extended Data Figure 7. Evaluating the influence of sample size on the accuracy of predicting ASD status at different ages. a,** Random samples were taken from the aggregate dataset of ASV's given the most resolved taxonomic assignment and trained for ASD classification using a GBC. Plotted are the obtained AUC's by sample size plotted with a non-linear fit and 95% confidence intervals. **b,** Heatmap of sample sizes when sub setting the aggregate dataset at various age groups. **c,** Study factors included as technical effects for batch correction, and the resulting AUC's for predicting ASD status with GBC models are reported. Data producing the greatest AUC after batch correction, along with the aggregated data containing no batch correction, and the data harboring the smallest AUC after batch correction are reported alongside their respective AUC's.

### Supplementary Figures

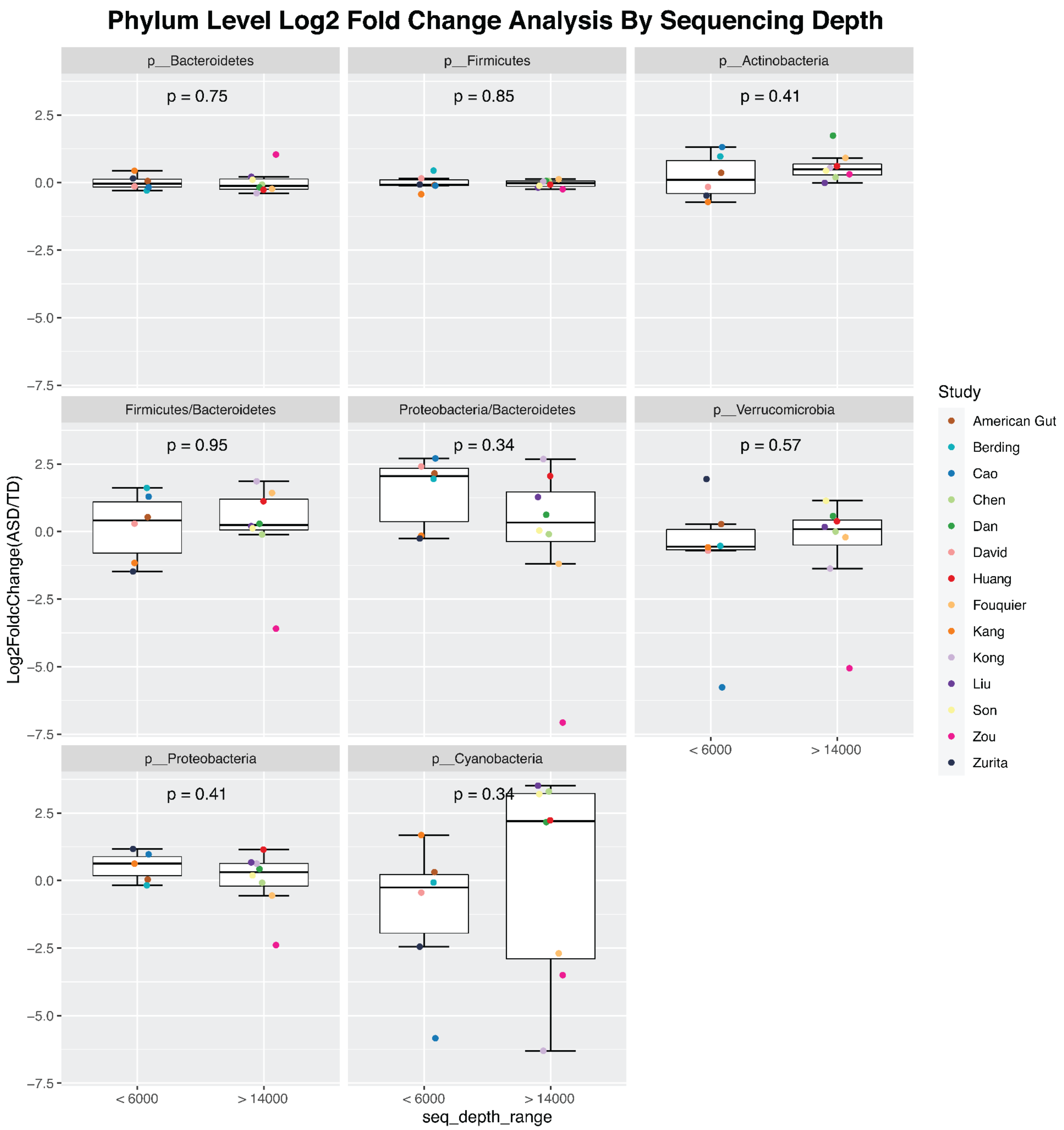


**Supplementary Figure 1. Effect of sequencing depth on ASD/NT phylum abundance.** Boxplots of the log2 fold change in ASD/controls for cohorts, processed individually, with a minimum sequencing depth of 6,000 (N = 6) compared to those greater than 14,000 (N = 8) for the phyla present in each cohort. P values were calculated from Wilcoxon two-tailed t-tests unadjusted for multiple hypothesis testing.

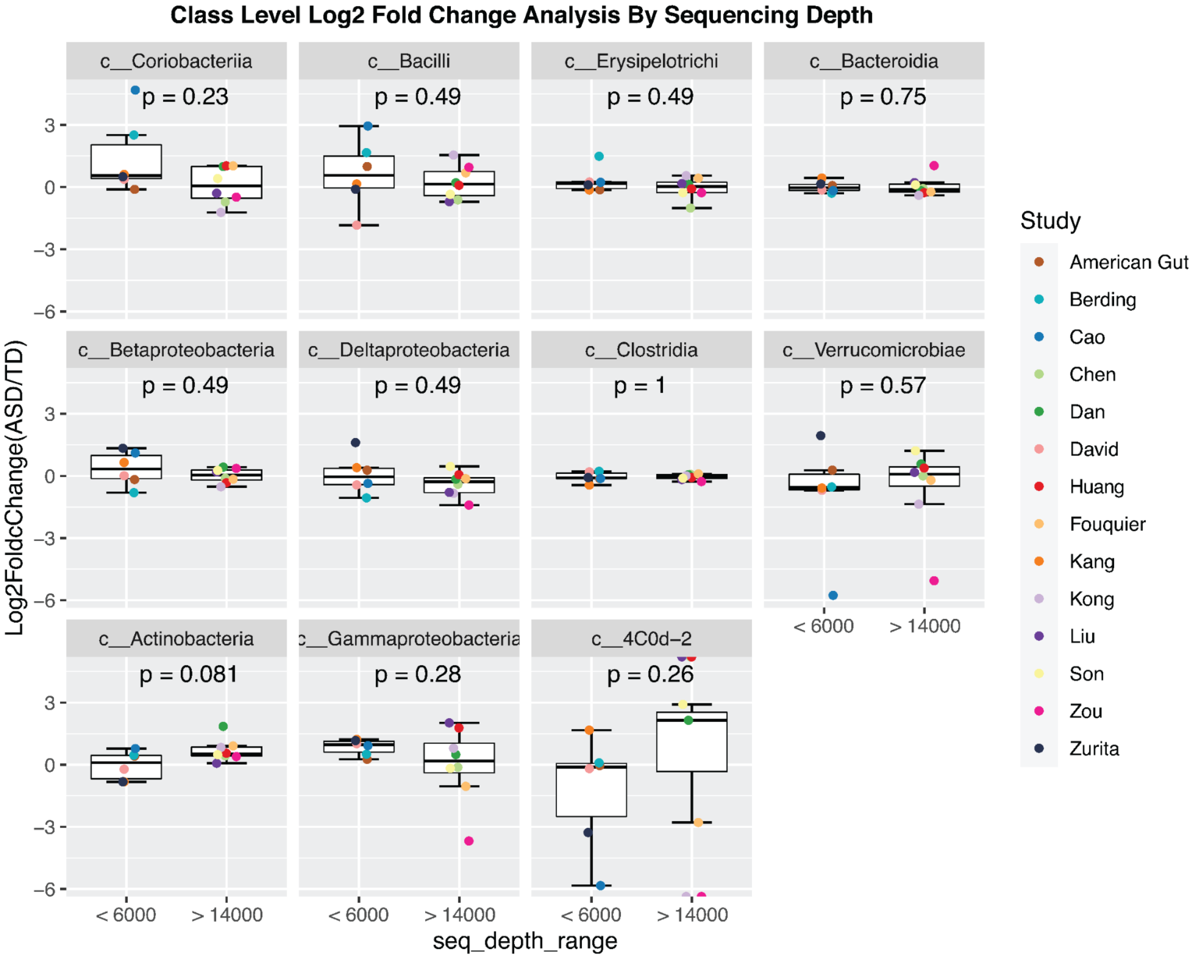


**Supplementary Figure 2. Effect of sequencing depth on ASD/NT class abundance.** Boxplots of the log2 fold change in ASD/controls for cohorts, processed individually, with a minimum sequencing depth of 6,000 (N = 6) compared to those greater than 14,000 (N = 8) for the classes present in each cohort. P values were calculated from Wilcoxon two-tailed t-tests unadjusted for multiple hypothesis testing.

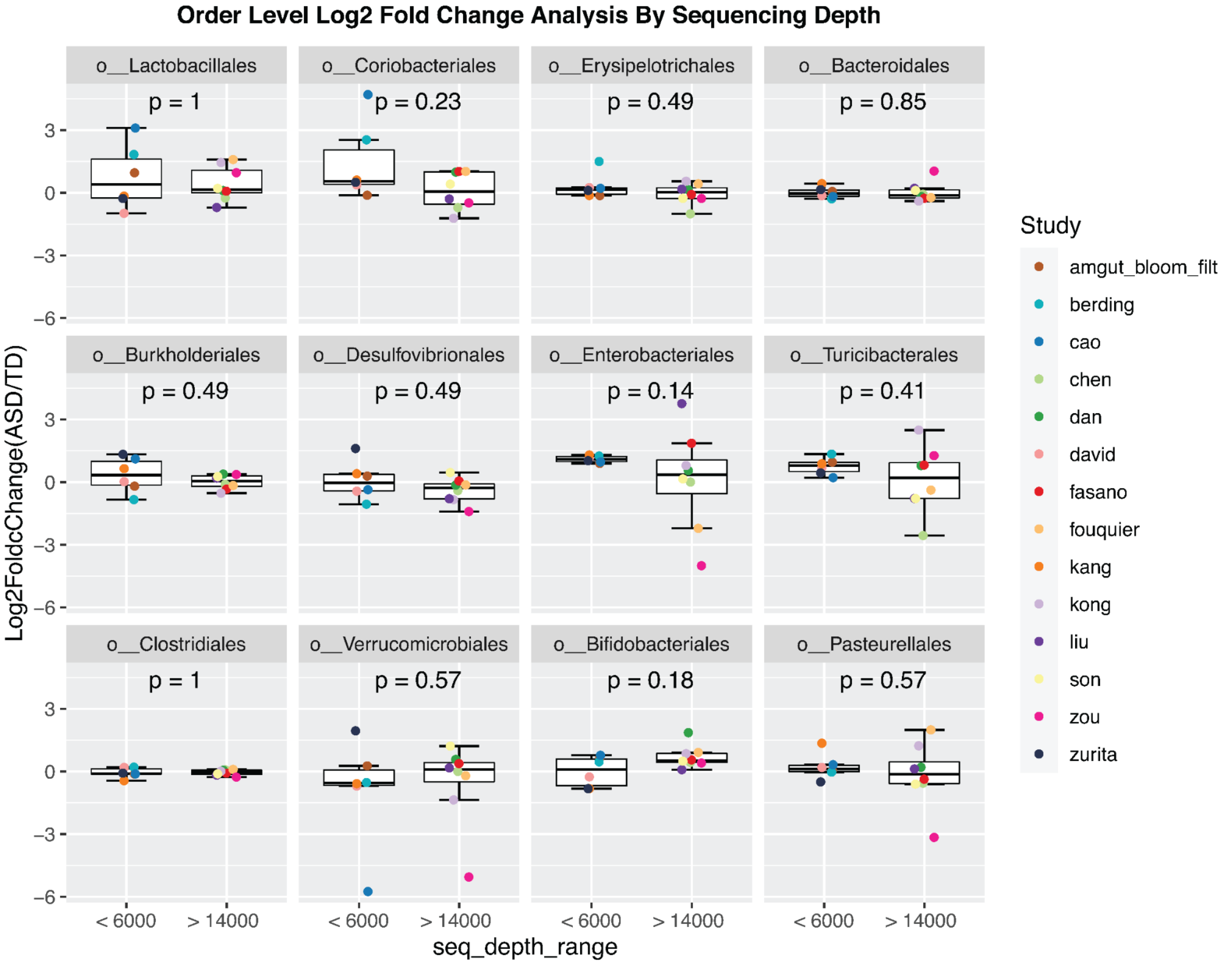


**Supplementary Figure 3. Effect of sequencing depth on ASD/NT order abundance.** Boxplots of the log2 fold change in ASD/controls for cohorts, processed individually, with a minimum sequencing depth of 6,000 (N = 6) compared to those greater than 14,000 (N = 8) for the orders present in each cohort. P values were calculated from Wilcoxon two-tailed t-tests unadjusted for multiple hypothesis testing.

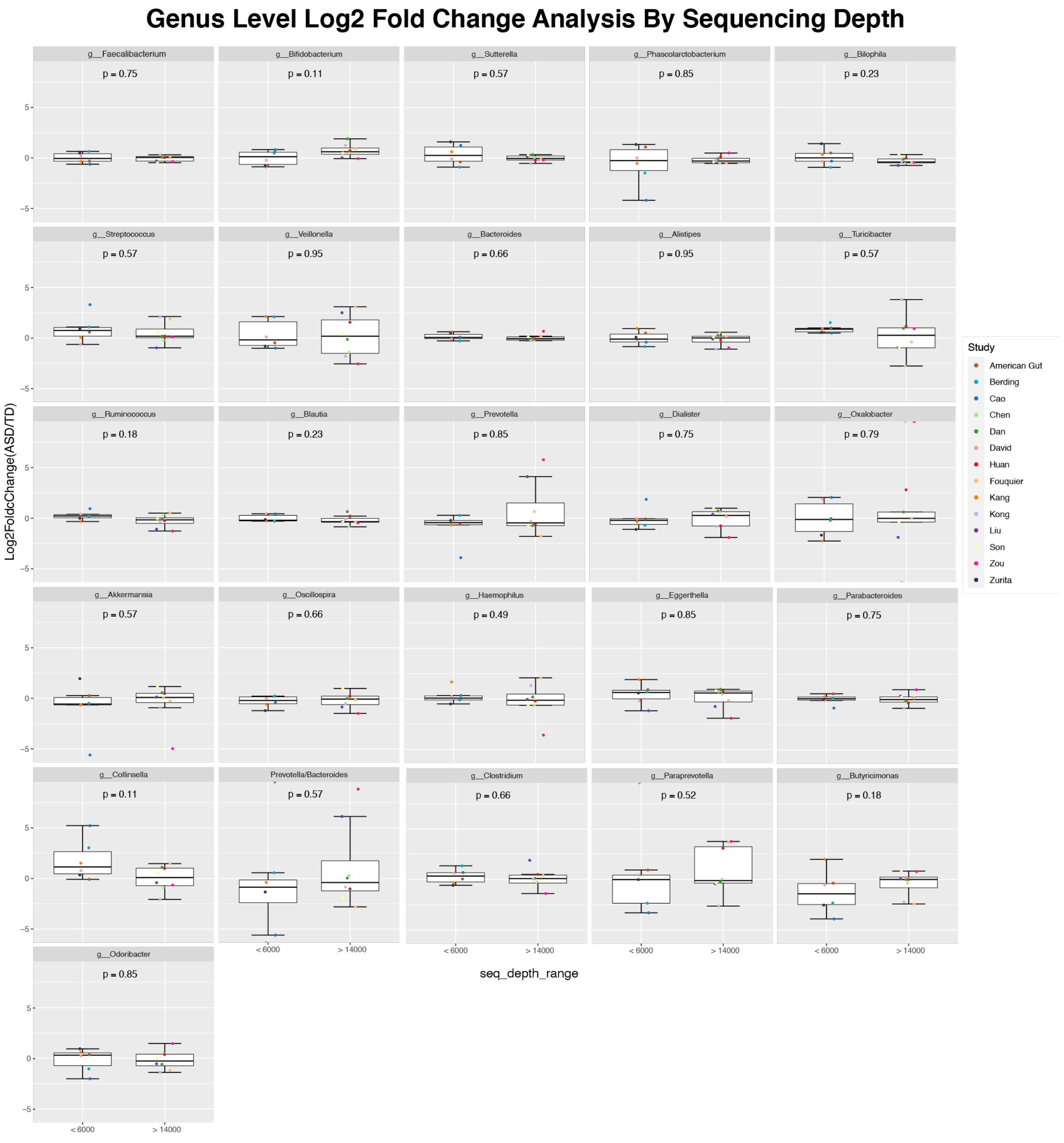


**Supplementary Figure 4. Effect of sequencing depth on ASD/NT genus abundance.** Boxplots of the log2 fold change in ASD/controls for cohorts, processed individually, with a minimum sequencing depth of 6,000 (N = 6) compared to those greater than 14,000 (N = 8) for the genera present in each cohort. P values were calculated from Wilcoxon two-tailed t-tests unadjusted for multiple hypothesis testing.

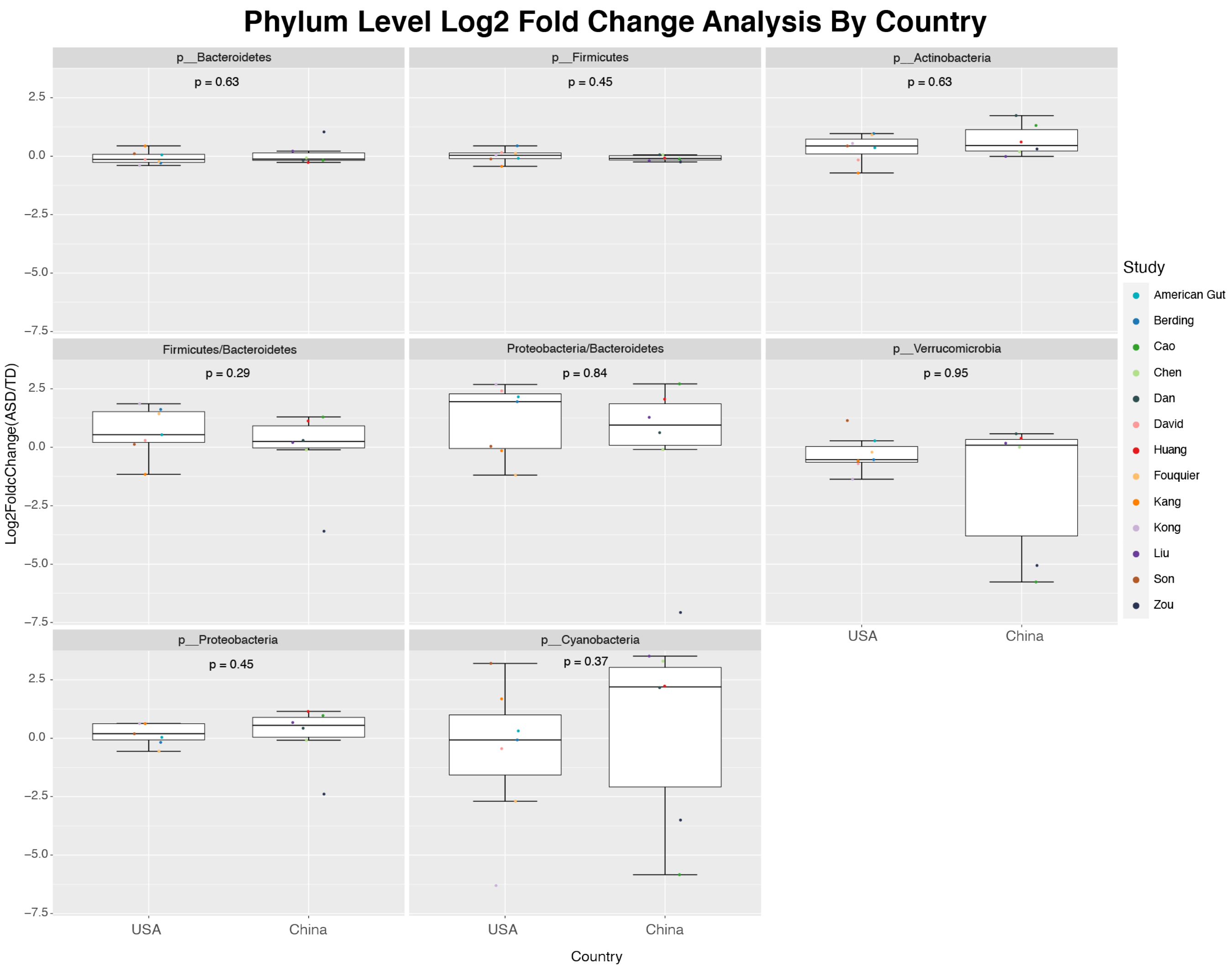


**Supplementary Figure 5. Effects of country on ASD/NT phylum abundance.** Boxplots of the log2 fold change in ASD/controls for individually processed cohorts are shown comparing data from the USA (N = 7) to data from China (N = 6) for phyla present in each cohort. P values were calculated from Wilcoxon two-tailed t-tests unadjusted for multiple hypothesis testing.

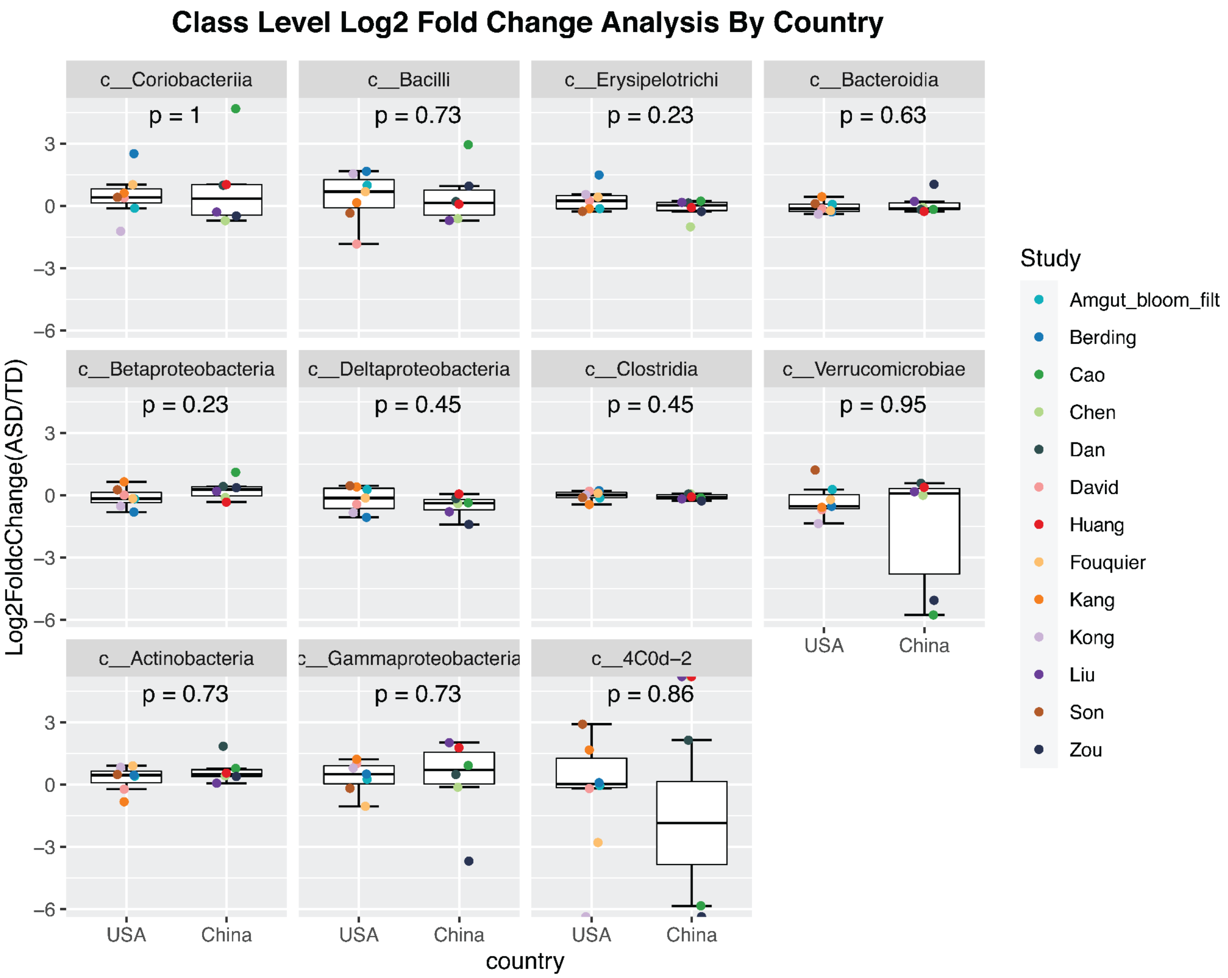


**Supplementary Figure 6. Effects of country on ASD/NT class abundance.** Boxplots of the log2 fold change in ASD/controls for individually processed cohorts are shown comparing data from the USA (N = 7) to data from China (N = 6) for classes present in each cohort. P values were calculated from Wilcoxon two-tailed t-tests unadjusted for multiple hypothesis testing.

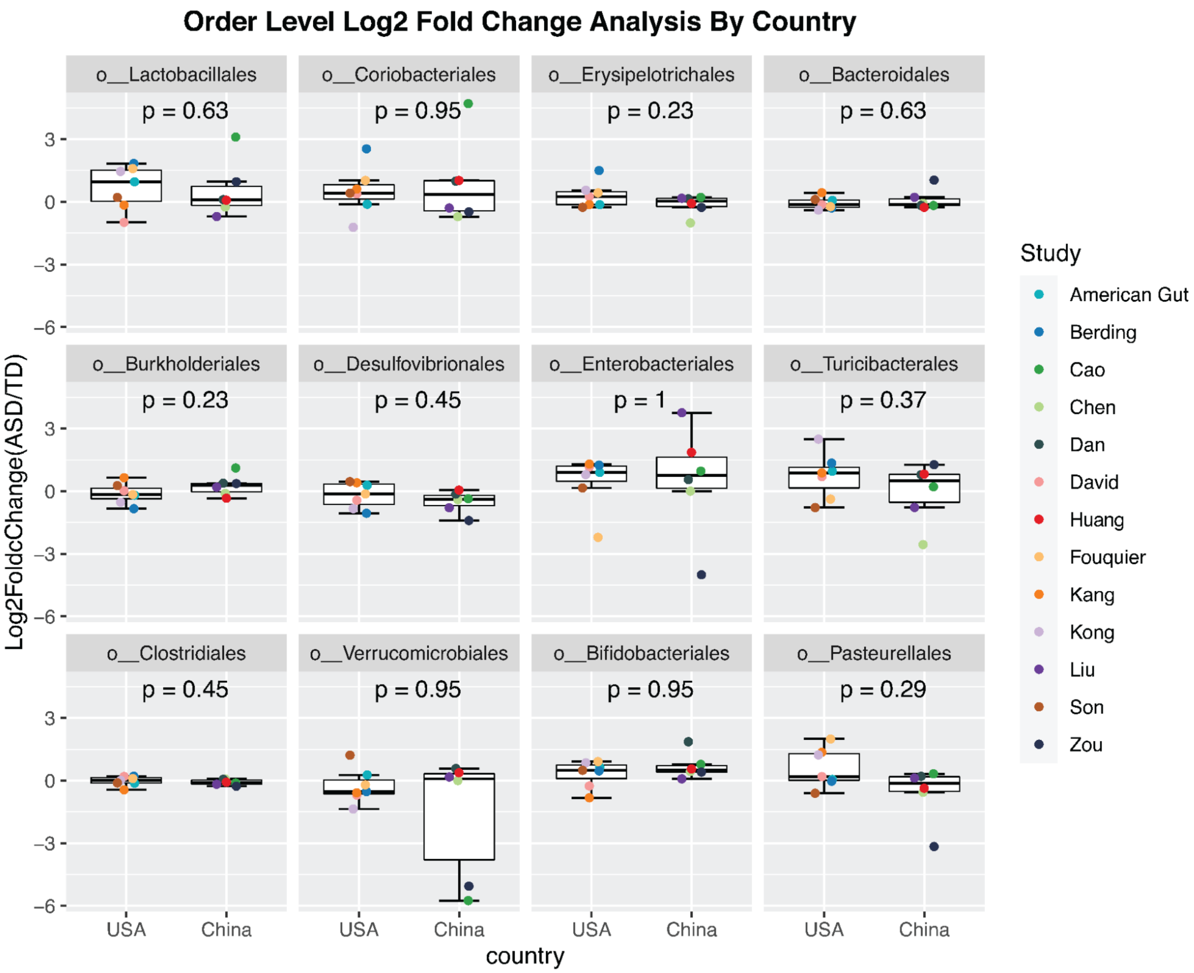


**Supplementary Figure 7. Effects of country on ASD/NT order abundance.** Boxplots of the log2 fold change in ASD/controls for individually processed cohorts are shown comparing data from the USA (N = 7) to data from China (N = 6) for orders present in each cohort. P values were calculated from Wilcoxon two-tailed t-tests unadjusted for multiple hypothesis testing.

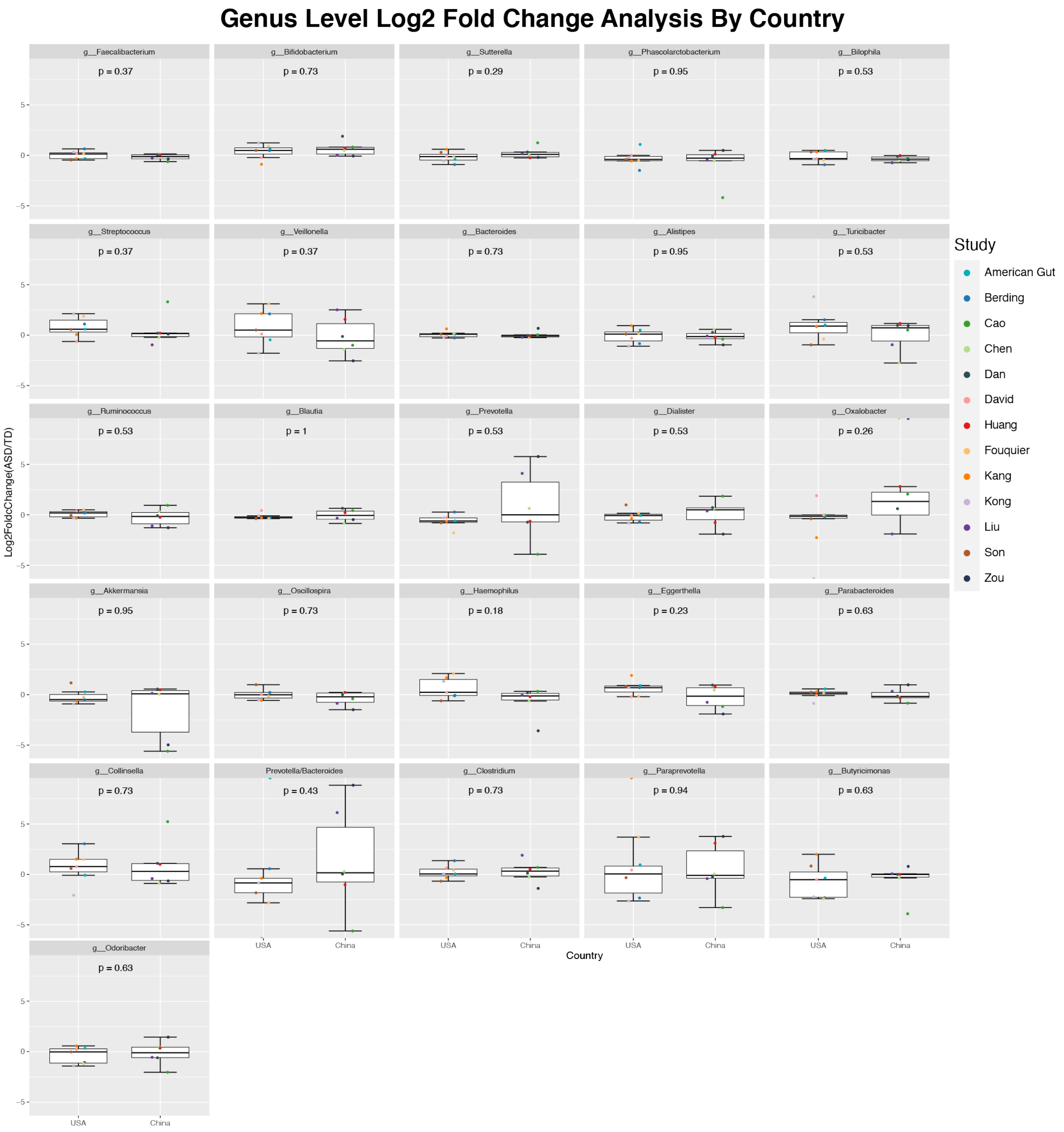


**Supplementary Figure 8. Effects of country on ASD/NT genus abundance.** Boxplots of the log2 fold change in ASD/controls for individually processed cohorts are shown comparing data from the USA (N = 7)

to data from China (N = 6) for genera present in each cohort. P values were calculated from Wilcoxon two-tailed t-tests unadjusted for multiple hypothesis testing.

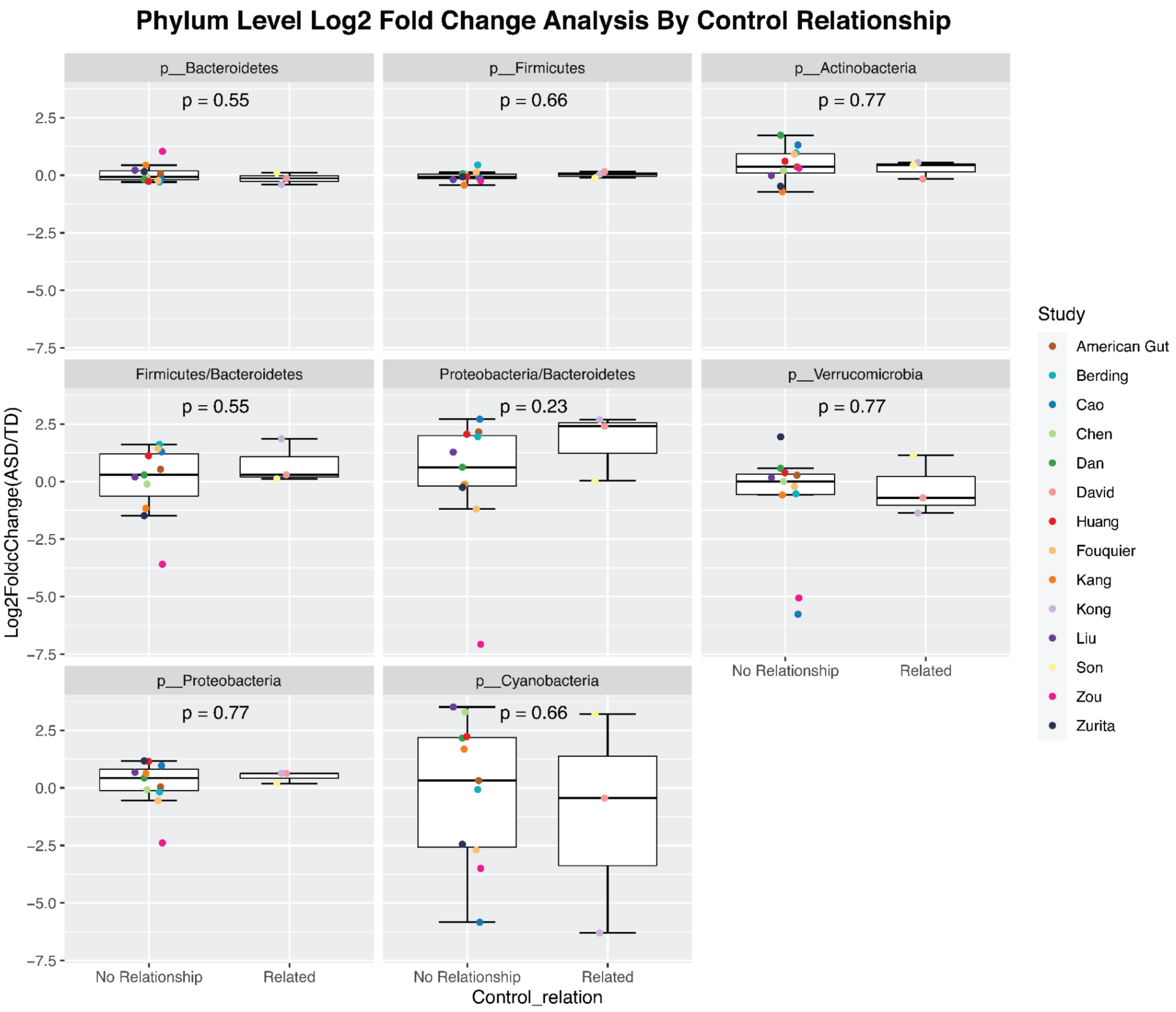


q

**Supplementary Figure 9. Effects of control type on ASD/NT phylum abundance.** Boxplots of the log2 fold change in ASD/controls for individually processed cohorts are shown comparing data collected using related controls (N = 3) to data collected from unrelated controls (N = 11) for phyla present in each cohort. P values were calculated from Wilcoxon two-tailed t-tests unadjusted for multiple hypothesis testing.

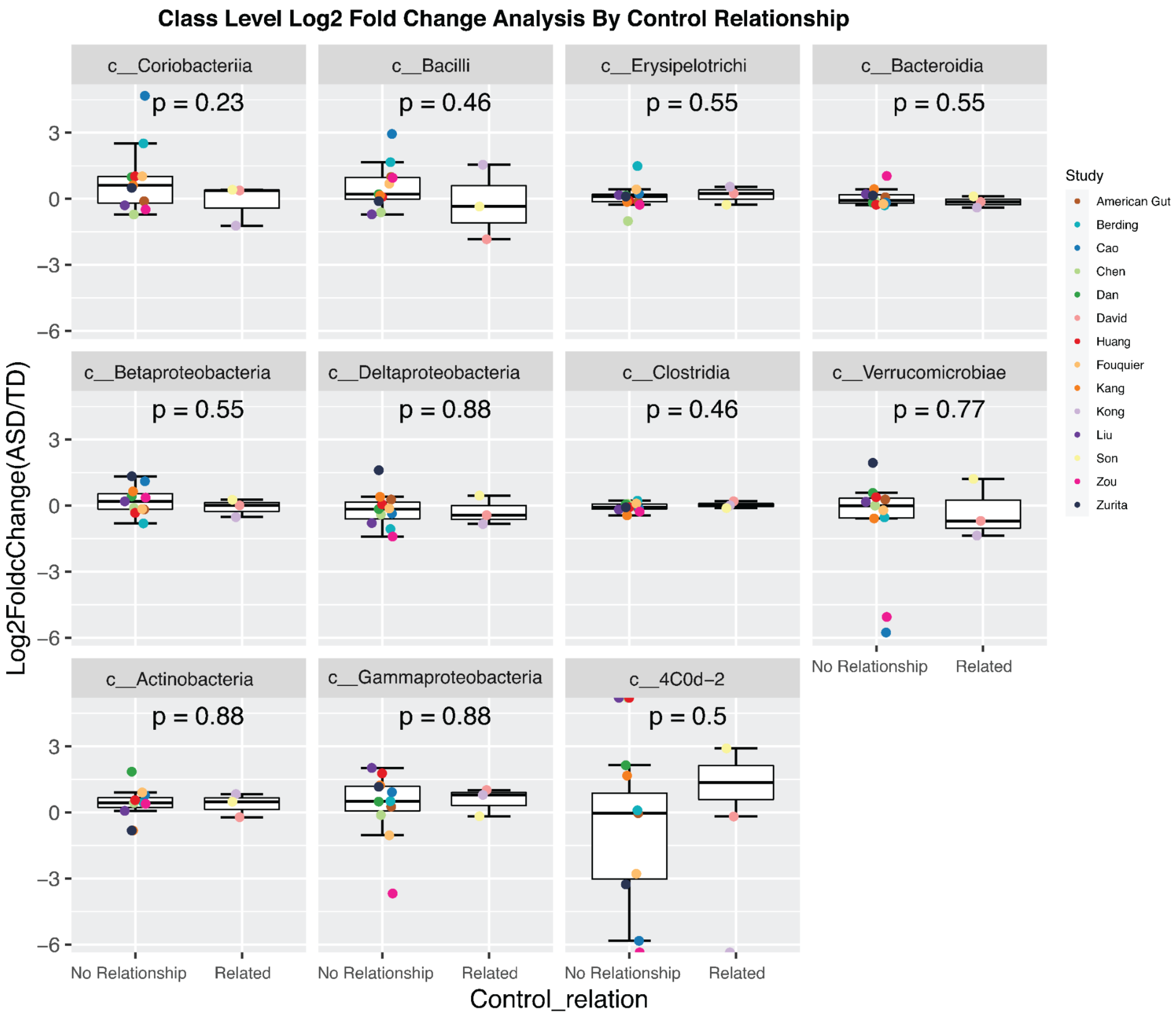


**Supplementary Figure 10. Effects of control type on ASD/NT class abundance.** Boxplots of the log2 fold change in ASD/controls for individually processed cohorts are shown comparing data collected using related controls (N = 3) to data collected from unrelated controls (N = 11) for classes present in each cohort. P values were calculated from Wilcoxon two-tailed t-tests unadjusted for multiple hypothesis testing.

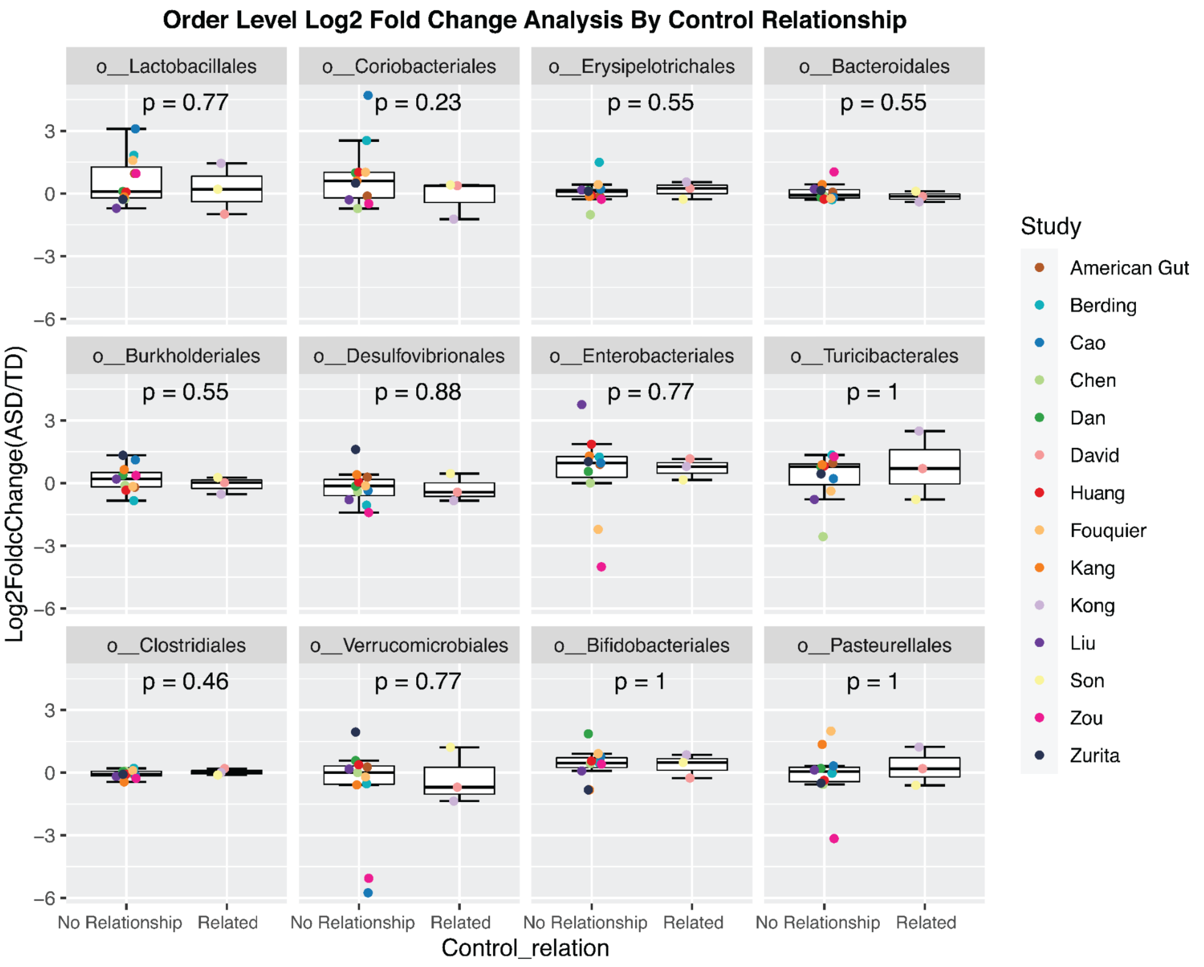


**Supplementary Figure 11. Effects of control type on ASD/NT order abundance.** Boxplots of the log2 fold change in ASD/controls for individually processed cohorts are shown comparing data collected using related controls (N = 3) to data collected from unrelated controls (N = 11) for orders present in each cohort. P values were calculated from Wilcoxon two-tailed t-tests unadjusted for multiple hypothesis testing.

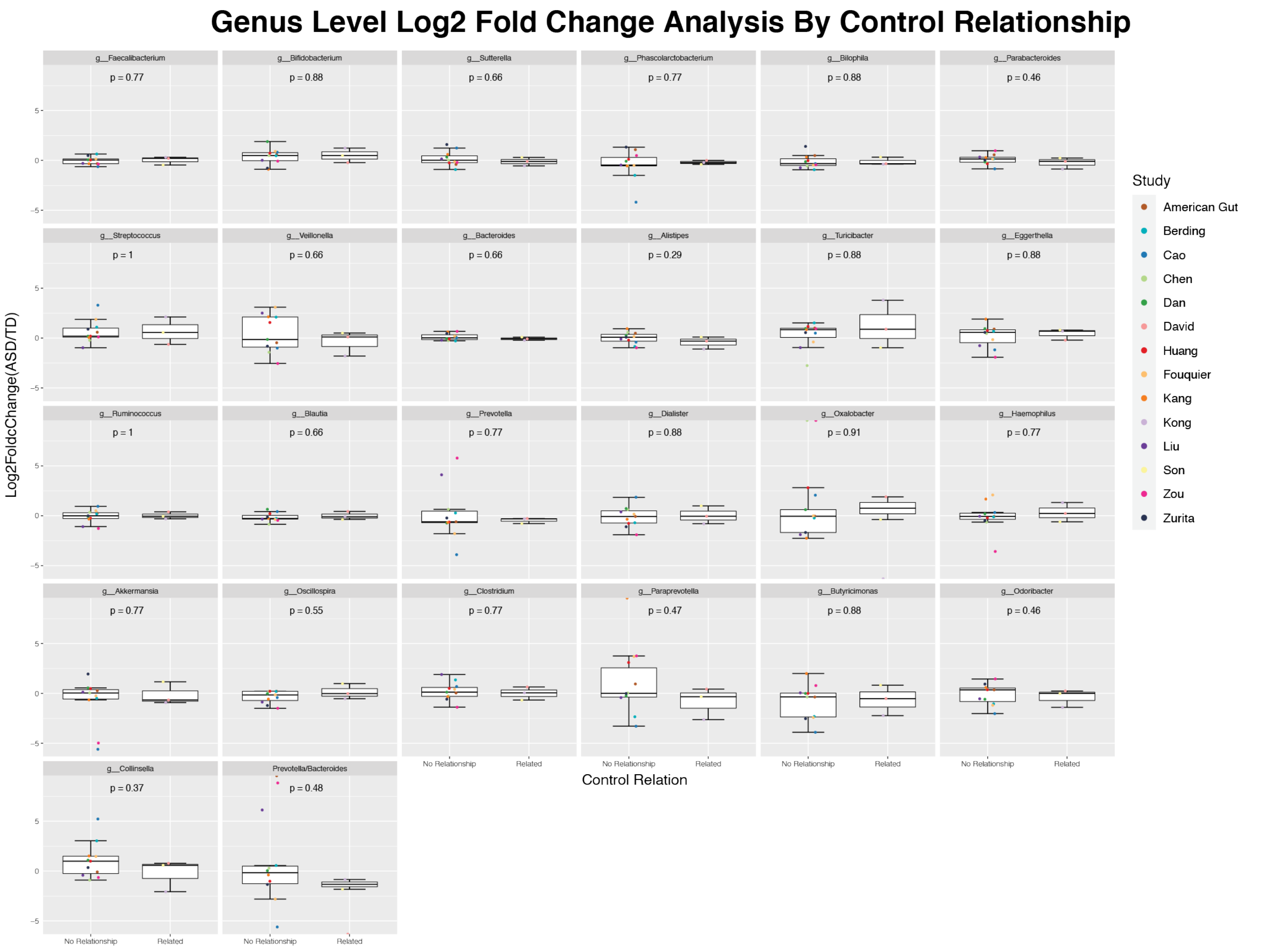


**Supplementary Figure 12. Effects of control type on ASD/NT genus abundance.** Boxplots of the log2 fold change in ASD/controls for individually processed cohorts are shown comparing data collected using related controls (N = 3) to data collected from unrelated controls (N = 11) for genera present in each cohort. P values were calculated from Wilcoxon two-tailed t-tests unadjusted for multiple hypothesis testing.
